# Moment-based approximations for the Wright-Fisher model of population dynamics under natural selection at two linked loci

**DOI:** 10.1101/2021.01.01.424882

**Authors:** Zhangyi He, Wenyang Lyu, Mark Beaumont, Feng Yu

## Abstract

Properly modelling genetic recombination and local linkage has been shown to bring significant improvement to the inference of natural selection from time series data of allele frequencies under a Wright-Fisher model. Existing approaches that can account for genetic recombination and local linkage are built on either the diffusion approximation or a moment-based approximation of the Wright-Fisher model. However, methods based on the diffusion approximation are likely to require much higher computational cost, whereas moment-based approximations may suffer from the distribution support issue: for example, the normal approximation can seriously affect computational accuracy. In the present work, we introduce two novel moment-based approximations of the Wright-Fisher model on a pair of linked loci, both subject to natural selection. Our key innovation is to extend existing methods to account for both the mean and (co)variance of the two-locus Wright-Fisher model with selection. We devise two approximation schemes, using a logistic normal distribution and a hierarchical beta distribution, respectively, by matching the first two moments of the Wright-Fisher model and the approximating model. As compared with the diffusion approximation, our approximations enable the approximate computation of the transition probabilities of the Wright-Fisher model at a far smaller computational cost. We can also avoid the distribution support issue found in the normal approximation.

## 1. Introduction

A central issue in evolutionary and population genetics is to understand the relative importance of genetic drift and natural selection as the mechanisms for maintaining polymorphisms within populations while at the same time driving divergence between populations. Mathematical descriptions of population dynamics under the effects of various demographic and evolutionary forces such as genetic drift and natural selection can be modelled through the Wright-Fisher model, introduced by Fisher (1922) and Wright (1931). To date, the Wright-Fisher model forms the basis of most theoretical and applied research in evolutionary and population genetics, *e.g.*, Kimura’s work on fixation probabilities (Kimura, 1955), Ewens’ sampling formula (Ewens, 1972) and Kingman’s coalescent (Kingman, 1982). Moreover, the increased availability of time series genetic data in recent years makes it possible to accurately infer demographic and genetic properties of observed populations, especially detection and estimation of natural selection. For this reason, the Wright-Fisher model is also a popular choice as the underlying mathematical model on which statistical techniques for inferring natural selection are built (see Tataru et al., 2017, for a review of such statistical inference techniques using allele frequency data).

The most important ingredient for carrying out statistical inference of natural selection from time series data of allele frequencies is to compute, conditioning on the initial allele frequency, the probability distribution of the allele frequency at each given time point. However, to our knowledge, there is no existing general tractable analytical form for the probability distribution of the allele frequency under the Wright-Fisher model. Numerical evaluation of the probability distribution of the allele frequency is theoretically feasible but computationally prohibitive for large population sizes and evolutionary timescales. Most existing approaches are therefore built on either the diffusion approximation of the Wright-Fisher model (*e.g.*, Bollback et al., 2008; Malaspinas et al., 2012; Steinrücken et al., 2014; Schraiber et al., 2016; Ferrer-Admetlla et al., 2016; He et al., 2020b,c) or moment-based approximations of the Wright-Fisher model (*e.g.*, Lacerda & Seoighe, 2014; Terhorst et al., 2015; Paris et al., 2019). These approximations enable efficient integration over all possible allele frequency trajectories of the underlying population, thereby allowing the likelihood computation to be completed in a reasonable amount of time.

Recent advances in sequencing techniques have revolutionised the collection of time series genomic data, with increased volume and improved quality (*e.g.*, Mathieson et al., 2015; Fages et al., 2019; Papkou et al., 2019; Barghi et al., 2019). This provides an opportunity for studying natural selection while accounting for the process of genetic recombination and the information of local linkage. Properly modelling genetic recombination and local linkage has been shown to significantly improve the inference of natural selection from genetic time series (see He et al., 2020b). However, with the exception of Terhorst et al. (2015) and He et al. (2020b), all existing Wright-Fisher model based methods for estimating selection coefficients from temporally spaced genetic data are limited to either a single locus (*e.g.*, Bollback et al., 2008; Malaspinas et al., 2012; Steinrücken et al., 2014; Lacerda & Seoighe, 2014; Schraiber et al., 2016; He et al., 2020c) or multiple independent loci (*e.g.*, Ferrer-Admetlla et al., 2016; Paris et al., 2019), *i.e.*, genetic recombination and local linkage are ignored in these approaches.

Terhorst et al. (2015), one of the exceptions amongst these methods, extended the moment-based approximation of the Wright-Fisher model introduced by Feder et al. (2014) to the case of natural selection acting on multiple linked loci, where the Wright-Fisher model was approximated through a deterministic path with added Gaussian noises. Their Gaussian approximation works well for many applications when modelling the allele frequencies with intermediate values. However, as soon as the allele frequencies approach their boundaries 0 or 1 (*i.e.*, allele loss or fixation), the Wright-Fisher model will be poorly approximated owing to the infinite support of the Gaussian distribution that will result in a leak of probability mass into unphysical values of the allele frequency that are smaller than 0 or larger than 1. This issue becomes more problematic in the inference of natural selection as natural selection is expected to rapidly drive the allele frequencies towards the boundaries. Most recently, He et al. (2020b) developed a Bayesian statistical framework for inferring natural selection at two linked loci while explicitly modelling genetic recombination and local linkage. Their approach was built upon the diffusion approximation of the Wright-Fisher model for a population evolving under natural selection at a pair of linked loci, represented in the stochastic differential equation form introduced by He et al. (2020a). The computation of the transition probability density function by numerically solving the resulting Kolmogorov backward equation (or its adjoint) is computationally challenging and prohibitively expensive. Their posterior computation was therefore carried out with the particle marginal Metropolis-Hastings algorithm developed by Andrieu et al. (2010), which avoids the computation of the transition probability density function but is still time-consuming.

In this work, we introduce two novel moment-based approximations for the Wright-Fisher model of population dynamics subject to natural selection at two linked loci. The main idea is to pick from a parametric family of probability distributions that matches the first two moments to those of the Wright-Fisher model. This is along the line of Cavalli-Sforza & Edwards (1967). To achieve the first two moments of the two-locus Wright-Fisher model with selection, we extend the moment approximations introduced by Lacerda & Seoighe (2014) and Paris et al. (2019), respectively. To avoid the distribution support issue found in the Gaussian approximation of Terhorst et al. (2015), we choose two different parametric families: logistic normal distributions and hierarchical beta distributions. As compared with the diffusion approximation of He et al. (2020b), our moment-based approximations enable us to compute the transition probabilities of the Wright-Fisher model at a much smaller computational cost. Our moment-based approximations are therefore expected to improve the performance of the existing statistical frameworks for the inference of natural selection from time series allele frequency data.

## 2. Wright-Fisher model

We consider a population of *N* randomly mating diploid individuals, consisting of a pair of linked loci, labelled 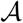 and 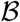, evolving under natural selection with discrete and nonoverlapping generations. Suppose that loci 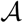 and 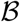 are located on the same chromosome with recombination rate *r* ∈ [0, 0.5] between them (*i.e.*, the rate that a recombinant gamete is produced at meiosis). At each locus, we assume that there are two possible allele types, labelled 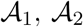 and 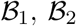, respectively, which gives rise to four possible haplotypes 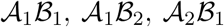 and 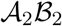, called haplotypes 1, 2, 3 and 4, respectively. Here we attach the symbols 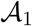 and 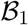 to the mutant alleles, which are assumed to arise only once in the population. The symbols 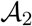 and 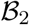 are the ancestral alleles, which are assumed to originally exist in the population.

We incorporate viability selection into population dynamics and let *w*_*ij*_ denote the viability of an individual with the genotype formed by haplotypes *i* and *j*. In addition to positing absence of sex effects, *i.e.*, *w*_*ij*_ = *w*_*ji*_, we assume that the viability of the genotype at the two loci is determined multiplicatively from the viabilities at individual loci and is fixed over time. More specifically, the viabilities of the three possible genotypes at each locus, *i.e.*, genotypes 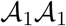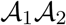 and 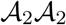 at a given locus 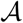, are taken to be 1, 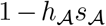 and 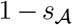, respectively, where 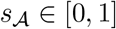 is the selection coefficient and 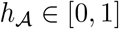 is the dominance parameter. For example, the viability of an individual carrying the 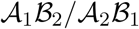 genotype is 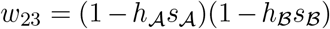.

Let *X*_*i*_(*k*) denote the frequency of gamete *i* in a population of *N* individuals in generation *k* ∈ ℕ for *i* = 1, 2, 3, 4, and ***X***(*k*) be the vector of the frequencies of the four possible gametes. In the Wright-Fisher model, the population size *N* is assumed to be fixed over time, and at each generation gametes are randomly sampled from an effectively infinite gamete pool reflecting the parental gamete frequencies. Under multinomial sampling, we have

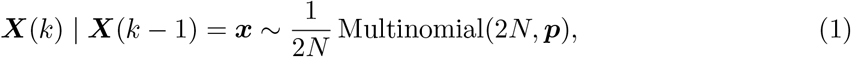

where ***p*** is the vector of the parental gamete frequencies of an effectively infinite population in generation *k* − 1 and can be written down as a function of the gamete frequencies ***x***,

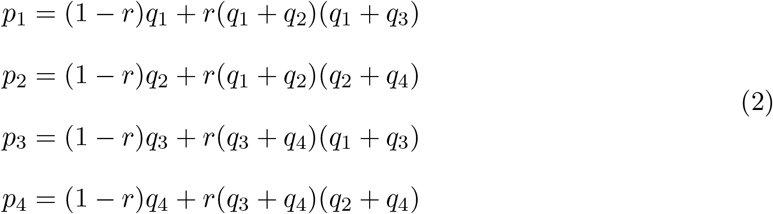

with

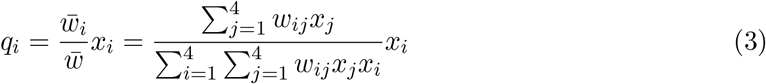

for *i* = 1, 2, 3, 4. In Eq. (3), the numerator 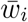 is the marginal fitness of haplotype *i* and the denominator 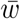 is the mean fitness.

We define the two-locus Wright-Fisher model with selection to be the stochastic process ***X*** = {***X***(*k*), *k* ∈ ℕ} evolving in the state space 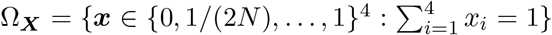 with the multinomial one-step transition probabilities as specified in Eqs. (1)–(3).

## 3. Moment-based approximations

Even though the one-step transition probabilities of the Wright-Fisher model, specified in Eqs. (1)–(3), can be written down, the *k*-step transition probabilities are intractable for general *k*. Numerical evaluation of the *k*-step transition probabilities is theoretically feasible but computationally prohibitive for large population sizes and evolutionary timescales. For this reason, approximations to the transition probabilities in the Wright-Fisher model based on parametric families have been developed, which could be traced back to Cavalli-Sforza & Edwards (1967). However, existing moment-based approximations either suffer from the distribution support issue (*e.g.*, Lacerda & Seoighe, 2014; Terhorst et al., 2015) or only work for the case of natural selection acting on a single locus (*e.g.*, Lacerda & Seoighe, 2014; Paris et al., 2019). To address these issues, we introduce two moment-based approximations for the two-locus Wright-Fisher model with selection. We first derive approximations for the mean and (co)variance, and then, to approximate the transition probabilities, we describe a logistic normal approximation and a hierarchical beta approximation.

### 3.1. Mean and (co)variance approximations

The main idea of the moment-based approximation is to approximate the transition probabilities of the Wright-Fisher model with a continuous probability distribution chosen from a parametric family, so that the first two moments are matched. This approach was first extended by Lacerda & Seoighe (2014) to incorporate natural selection and using normal distributions, applied to the inference of natural selection from time series data of allele frequencies. Its prerequisite is to compute the mean and (co)variance of the Wright-Fisher model with selection at any given time point. Terhorst et al. (2015) proposed an iteration formula to approximate the first two moments of the Wright-Fisher model, which is the only method applicable to the case of natural selection acting on multiple linked loci. We extend other two moment approximations developed in Lacerda & Seoighe (2014) and Paris et al. (2019) to the two-locus Wright-Fisher model with selection.

From Eq. (1), given the gamete frequencies ***X***(*k* − 1) = ***x***, we can write down the mean and (co)variance of the gamete frequencies ***X***(*k*) as

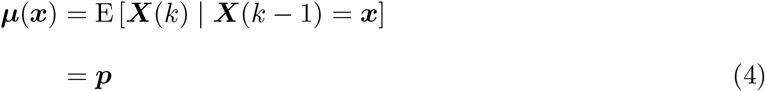

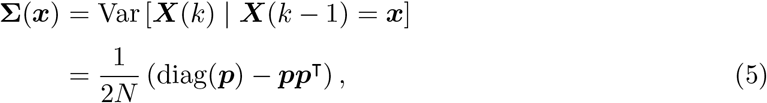

respectively, where diag(***p***) is the operator that constructs a diagonal matrix and places the elements of the sampling probability vector ***p*** on the main diagonal.

Let ***μ***_*k*_ and **Σ**_*k*_ denote the mean and (co)variance of the gamete frequencies ***X***(*k*) given the gamete frequencies ***X***(0) for *k* ∈ ℕ^+^, respectively. Using the law of total mean and (co)variance, we have

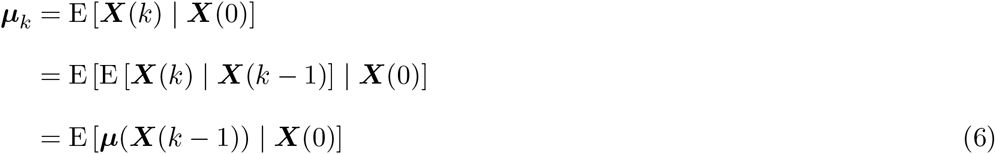

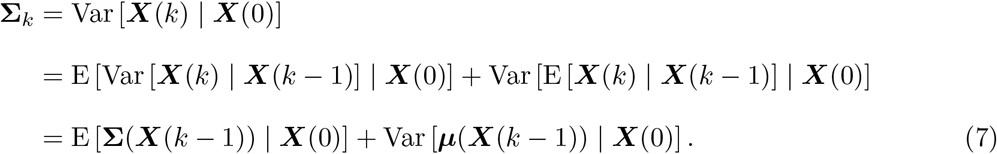

Following Lacerda & Seoighe (2014), we apply the delta method with the first-order Taylor expansion of the term ***μ***(***X***(*k* − 1)) about the mean ***μ***_k−1_. The mean ***μ***_*k*_ and the (co)variance **Σ**_*k*_ can then be approximated as

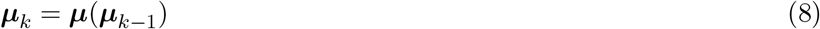

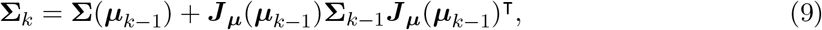

where ***J***_***μ***_(***x***) is the Jacobian matrix defined over the function ***μ***(***x***) in Eq. (4). By substituting Eqs. (2) and (3) into Eq. (4) and taking the derivative with respect to the gamete frequencies ***x***, we have

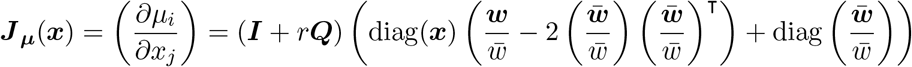

with

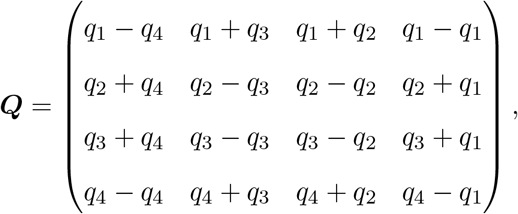

where ***w*** = (*w*_*ij*_) is the viability matrix for all genotypes, and 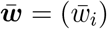 is the marginal viability vector for all haplotypes.

Alternatively, following Paris et al. (2019), we substitute Eqs. (4) and (5) into Eq. (7). Then the (co)variance **Σ**_*k*_ can be expressed as

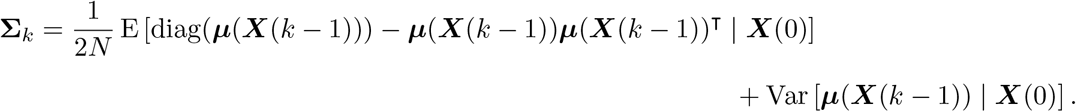

With the first-order Taylor expansion of the term ***μ***(***X***(*k* − 1)) about the mean ***μ***_*k*−1_, we can approximate the mean ***μ***_*k*_ and the (co)variance **Σ**_*k*_ as

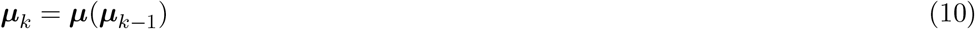

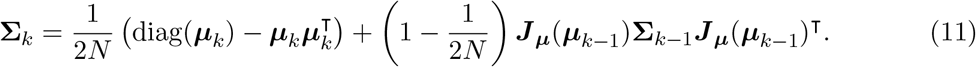

With the second-order Taylor expansion of the term ***μ***(***X***(*k* − 1)) about the mean ***μ***_*k*−1_, we can approximate the mean ***μ***_*k*_ and the (co)variance **Σ**_*k*_ as

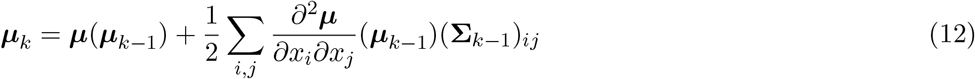

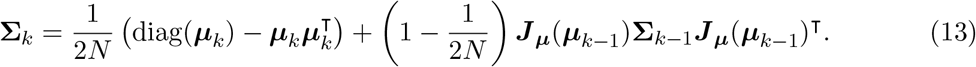

Using the iteration formula in Eqs. (8) and (9) or in Eqs. (10) and (11) or in Eqs. (12) and (13), we can achieve the approximations of the mean and (co)variance of the two-locus Wright-Fisher model with selection, ***μ***_*k*_ and **Σ**_*k*_, for *k* ∈ ℕ with initial values of ***μ***_0_ = E[***X***(0)] and **Σ**_0_ = Var[***X***(0)]. See the detailed derivation of the extension of the two moment approximations developed by Lacerda & Seoighe (2014) and Paris et al. (2019) for the two-locus Wright-Fisher model with selection in Appendix A, where the detailed derivation of the moment approximation of Terhorst et al. (2015) can also be found.

### 3.2. Logistic normal approximation

With the approximations for the first two moments of the two-locus Wright-Fisher model with selection, we now set out to find distributions to approximate the true transition probabilities. We first consider the family of normal distributions, which has already been successfully used in population genetics studies (*e.g.*, Nicholson et al., 2002; Pickrell & Pritchard, 2012; Lacerda & Seoighe, 2014; Terhorst et al., 2015). We assume that the gamete frequencies ***X***(*k*) are approximately distributed according to

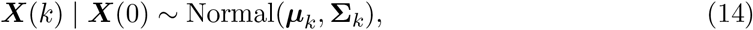

where ***μ***_*k*_ and **Σ**_*k*_ are the mean and (co)variance of the two-locus Wright-Fisher model with selection, respectively, which can be obtained by running the iteration formula in Eqs. (8) and (9) or in Eqs. (10) and (11) or in Eqs. (12) and (13) with the initial values ***μ***_0_ and **Σ**_0_.

The normal approximation is motivated by matching the mean and (co)variance of the normal distribution to those of the Wright-Fisher model. Unfortunately, the normal approximation in Eq. (14) is able to take values outside the state space Ω_***X***_, resulting in poor performance once gamete frequencies ***X***(*k*) approach the boundaries of the state space Ω_***X***_. The same distribution support issue can be found in Terhorst et al. (2015). In the single-locus case, to narrow the state space of the normal approximation, Nicholson et al. (2002) allocated the mass of the normal distribution outside the state space of the Wright-Fisher model to the discrete fixation probabilities at boundaries 0 and 1. As an alternative, Paris et al. (2019) simply rejected the values outside the state space of the Wright-Fisher model and rescaled the normal distribution. However, both these procedures lead to the mean and (co)variance of the resulting probability distribution no longer matching those of the Wright-Fisher model.

To tackle the distribution support issue of the normal approximation, we consider the logistic normal distribution with the support 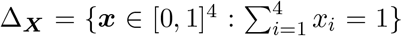. More specifically, we construct a transformation, denoted by ***φ***, to map the gamete frequencies ***X***(*k*) to the state space ℝ^3^:

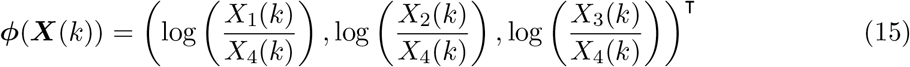

and introduce a stochastic process, denoted by ***Y*** = {***Y*** (*k*), *k* ∈ ℕ}, to approximate the transformed Wright-Fisher model ***φ***(***X***). More concretely, we let ***Y***(*k*) = (*Y*_1_(*k*), *Y*_2_(*k*), *Y*_3_(*k*))^T^ ∈ ℝ^3^ follow a normal distribution

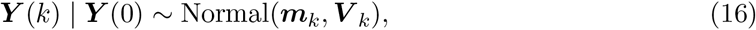

where ***m***_*k*_ and ***V***_*k*_ are the mean and (co)variance of the normal distribution. The unique inverse mapping given by

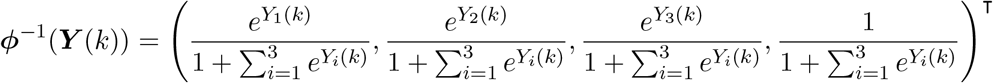

yields our moment-based approximation of the Wright-Fisher model ***X***, where the inverse mapping ***φ***^−1^ is commonly known as the additive logistic transformation. Given the initial gamete frequencies ***X***(0), our moment-based approximation ***φ***^−1^(***Y*** (*k*)) suggests that the gamete frequencies ***X***(*k*) approximately follows a logistic normal distribution

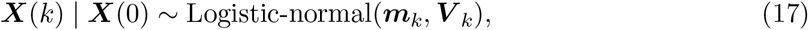

which we refer to as the logistic normal approximation of the two-locus Wright-Fisher model with selection.

Our task now is to find proper values for the mean and (co)variance of the normal distribution in Eq. (16) such that the logistic normal distribution in Eq. (17) matches its mean and (co)variance to those of the Wright-Fisher model ***X***. To the best of our knowledge, there is no analytical solution for the mean and (co)variance of the logistic normal distribution, but using the delta method with the first-order Taylor expansion of the term ***φ***(***X***(*k*)) about the mean ***μ***_***k***_, we can approximate the mean ***m***_*k*_ and the (co)variance ***V*** _*k*_ of the normal distribution with the mean ***μ***_*k*_ and the (co)variance **Σ**_*k*_ of the Wright-Fisher model ***X***,

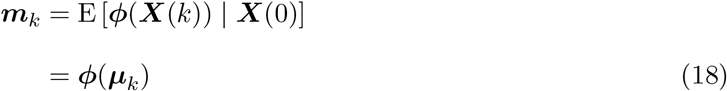

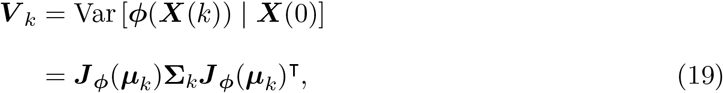

where

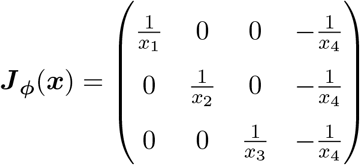

is the Jacobian matrix defined over the function ***φ***(***x***) in Eq. (15). Note that the mean ***μ***_*k*_ and the (co)variance **Σ**_*k*_ of the Wright-Fisher model ***X*** are readily available by running the iteration formula in Eqs. (8) and (9) or in Eqs. (10) and (11) or in Eqs. (12) and (13) with the initial values ***μ***_0_ and **Σ**_0_.

By substituting Eqs. (18) and (19) into Eq. (17), our logistic normal approximation of the two-locus Wright-Fisher model with selection can therefore be represented as

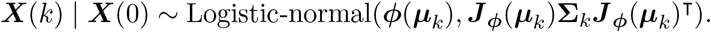

The transition probability density function of our logistic normal approximation can be explicitly written down by using the change-of-variable technique. Notice that the first two moments of our logistic normal approximation are no longer exactly matched to those of the Wright-Fisher model since the delta method is used to approximate the mean ***m***_*k*_ and the (co)variance ***V*** _*k*_ of the normal distribution in Eqs. (18) and (19) (see Figure 1 for an illustration). In Section 4, we will explore how such a discrepancy affects the performance of our logistic normal approximation through extensive simulations.

**Figure 1:**
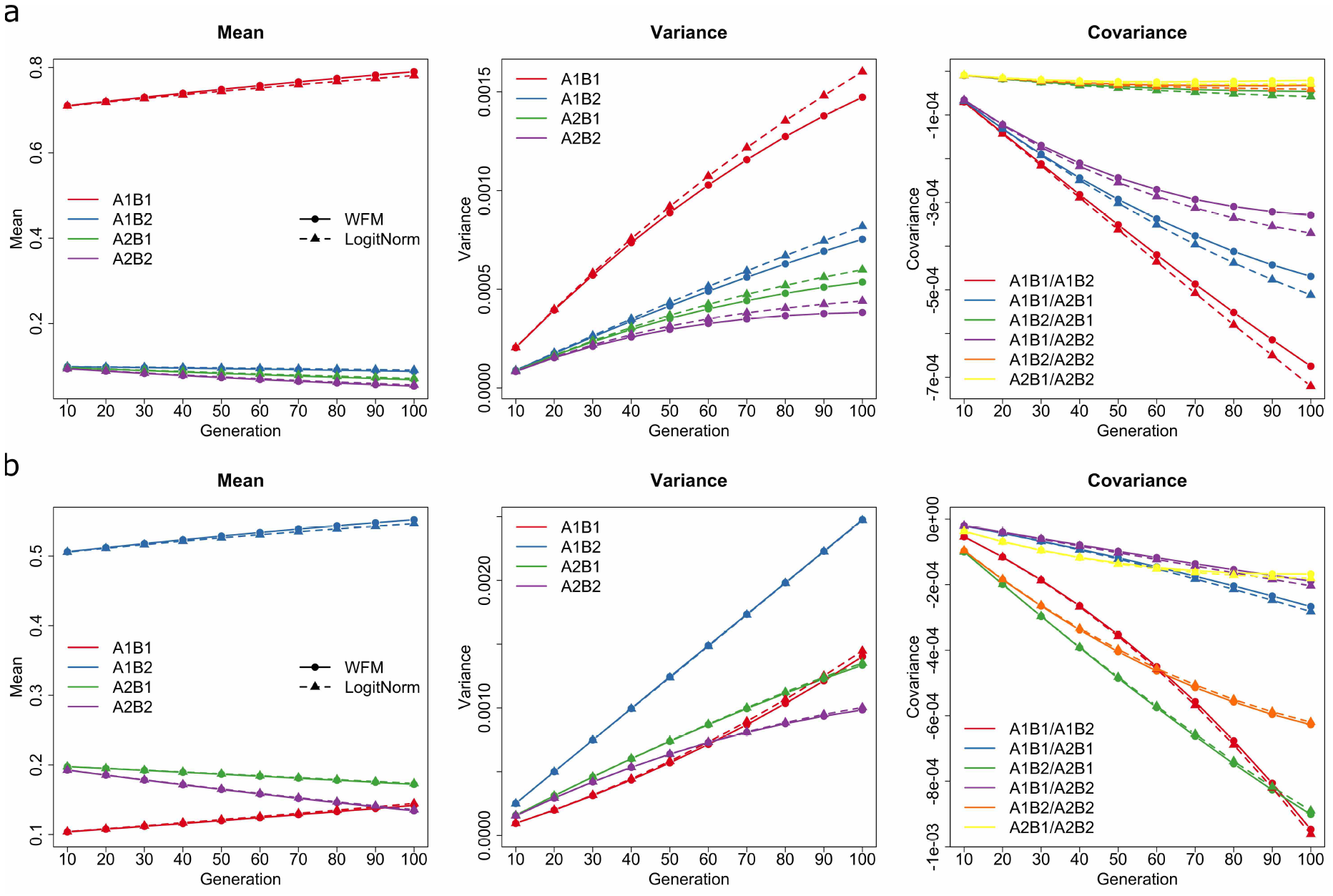
An illustration for the mean, variance and covariance of the Wright-Fisher model and our logistic normal approximation, where we set the selection coefficients 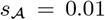 and 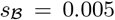, the dominance parameters 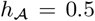 and 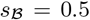, the recombination rate *r* = 0.00001 and the population size *N* = 5000. We pick the initial gamete frequencies (a) ***X***0 = (0.7, 0.1, 0.1, 0.1) and (b) ***X***_0_ = (0.1, 0.5, 0.2, 0.2), respectively. The first two moments of the Wright-Fisher model and the logistic normal approximation are estimated from 1,000,000 replicated simulations.

### 3.3. Hierarchical beta approximation

In addition to normal distributions, the beta family of distributions is an alternative candidate of the parametric family for the moment-based approximation of the Wright-Fisher model. This has been shown to deliver superior performance in comparison with normal distributions for single-locus problems (see Tataru et al., 2017; Paris et al., 2019) and successfully applied in a number of population genetics studies (*e.g.*, Sirén et al., 2011; Hui & Burt, 2015; Tataru et al., 2015; Paris et al., 2019). For two-locus problems, we need a multivariate generalisation of beta distributions, but unfortunately, the multivariate beta distribution, commonly known as the Dirichlet distribution, is not appropriate since its mean and (co)variance fail to fully match those of the two-locus Wright-Fisher model with selection. More specifically, the Dirichlet distribution of order four has four parameters, three of which are used to determine the mean of the Wright-Fisher model, leaving one free parameter to model the (co)variance. However, a single parameter is not enough to completely capture the (co)variance structure of the Wright-Fisher model.

Inspired by Hobolth & Siren (2016), we employ the hierarchical beta distribution with the support 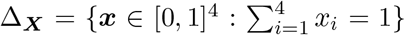, which is an alternative multivariate generalisation of the beta distribution. The main idea is to decompose the two-locus Wright-Fisher model with selection into two conditionally independent beta distributed levels. More specifically, we consider any three linearly independent gamete frequencies out of the four, which is illustrated below for *X*_1_(*k*), *X*_2_(*k*) and *X*_3_(*k*), and construct a transformation, denoted by ***ψ***, to map the gamete frequencies ***X***(*k*) to the state space [0, 1]^3^:

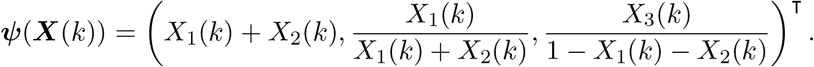

We introduce a stochastic process, denoted by ***Z*** = {***Z***(*k*), *k* ∈ ℕ}, to approximate the transformed Wright-Fisher model ***φ***(***X***). More concretely, let ***Z***(*k*) = (*Z*_1_(*k*), *Z*_2_(*k*), *Z*_3_(*k*))^T^ ∈ [0, 1]^3^ follow a hierarchical beta distribution

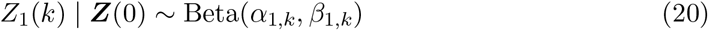

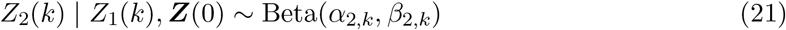

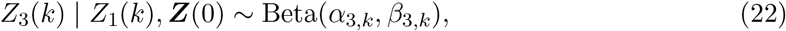

where *α*_*i,k*_ and *β*_*i,k*_ are the shape parameters of each beta distribution in Eqs. (20)–(22). Notice that *Z*_1_(*k*) is the frequency of the 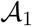 allele, *Z*_2_(*k*) is the frequency of the 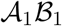 haplotype in the haplotypes with the 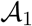 allele, and *Z*_3_(*k*) is the frequency of the 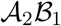 haplotype in the haplotypes with the 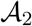 allele. The unique inverse mapping given by

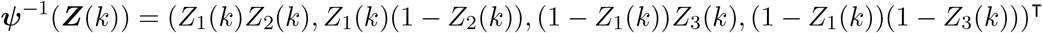

yields our moment-based approximation of the Wright-Fisher model ***X***. We call the procedure we just outlined the hierarchical beta approximation of the two-locus Wright-Fisher model with selection.

Our task now is to find proper values for the shape parameters *α*_*i,k*_ and *β*_*i,k*_ of each beta distribution in Eqs. (20)–(22) such that the probability distribution for our hierarchical beta approximation ***φ***^−1^(***Z***(*k*)) has its mean and (co)variance matching those of the Wright-Fisher model ***X***. The shape parameters of the beta distribution are related its mean and variance as follows:

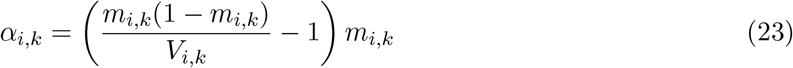

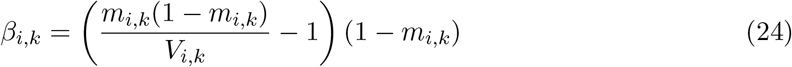

for *i* = 1, 2, 3, where *m*_*i,k*_ and *V*_*i,k*_ denote the mean and variance of each beta distribution in Eqs. (20)–(22). Our problem then reduces to the calculation of the mean *m*_*i,k*_ and the variance *V*_*i,k*_ of each beta distribution with the mean ***μ***_*k*_ and the (co)variance **Σ**_*k*_ of the Wright-Fisher model ***X***.

Using the law of total mean, we can now write down the mean of the frequency of the 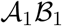 gamete under the Wright-Fisher model ***X*** in terms of the mean *m_i,k_* of each beta distribution as

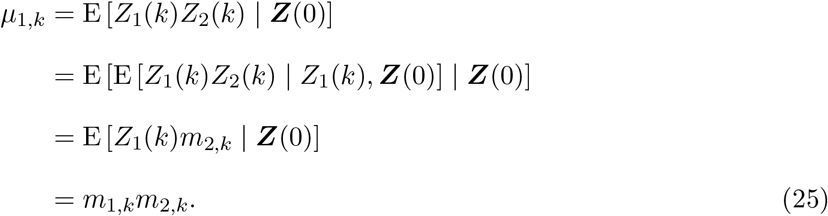

Similarly, the mean of the other three gamete frequencies can be written down as

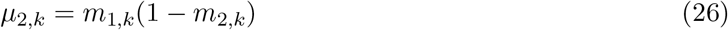

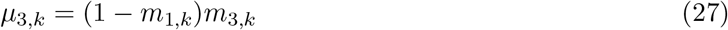

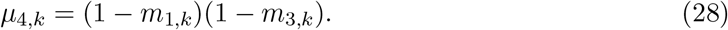

Solving the system of coupled equations given by Eqs. (25)–(27) yields

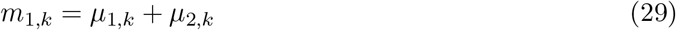

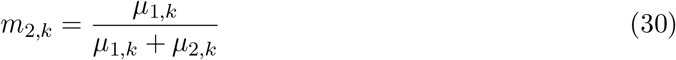

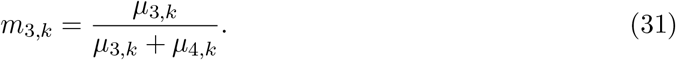

Note that the mean *m*_*i,k*_ given in Eqs. (29)–(31) also satisfies Eq. (28).

With the law of total variance, the variance of the frequency of the 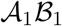 gamete under the Wright-Fisher model ***X*** can be represented in terms of the mean *m*_*i,k*_ and the variance *V*_*i,k*_ of each beta distribution as

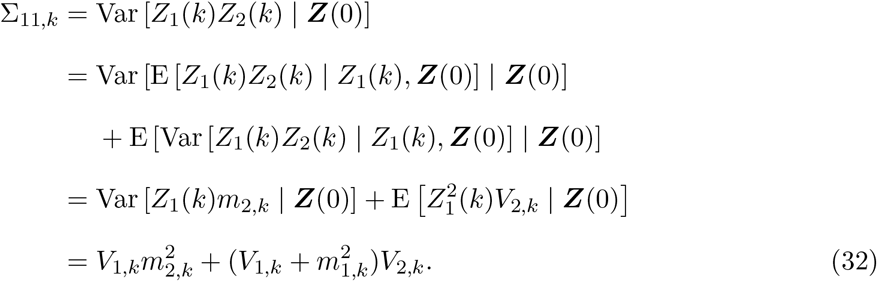

Similarly, the variance of the other three gamete frequencies can be expressed as

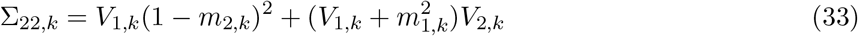

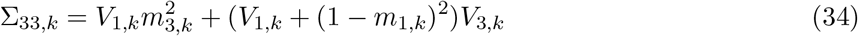

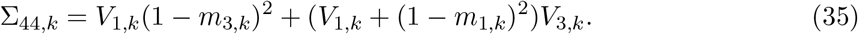

Solving the system of coupled equations given by Eqs. (32)–(34) yields

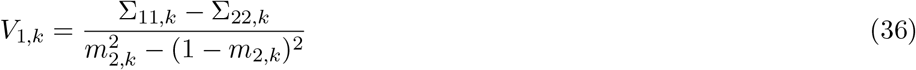

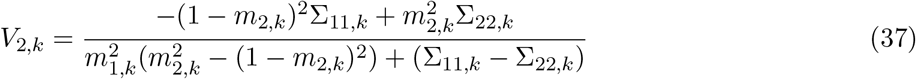

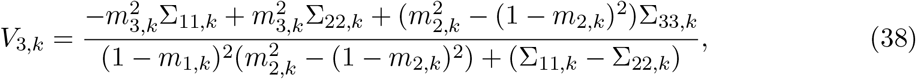

where *m*_*i,k*_ is the mean of each beta distribution given by Eqs. (29)–(31). Note that the variance *V*_*i,k*_ given in Eqs. (36)–(38) is no longer guaranteed to satisfy Eq. (35).

Recall that the mean ***μ***_*k*_ and the (co)variance **Σ**_*k*_ of the Wright-Fisher model ***X*** are readily available by running the iteration formula in Eqs. (8) and (9) or in Eqs. (10) and (11) or in Eqs. (12) and (13) with the initial values ***μ***_0_ and **Σ**_0_. Using Eqs. (20)–(22), we can therefore represent our hierarchical beta approximation of the two-locus Wright-Fisher model with selection as

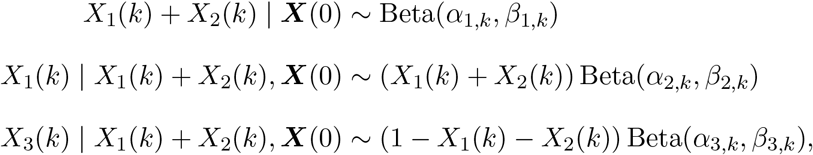

where the shape parameters *α_i,k_* and *β_i,k_* of each beta distribution can be obtained by substituting Eqs. (29)–(31) and (33)–(35) into Eqs. (23) and (24). The transition probability density function of our hierarchical beta approximation can be explicitly written down by the change-of-variable technique. Note that in our hierarchical beta approximation, the mean of our hierarchical beta approximation exactly matches those of the Wright-Fisher model, but only three of four variances are exactly matched (see Figure 2 for an illustration). This is because in our hierarchical beta approximation there is no extra free parameter left for fitting the one remaining variance and all covariance of the two-locus Wright-Fisher model with selection.

**Figure 2:**
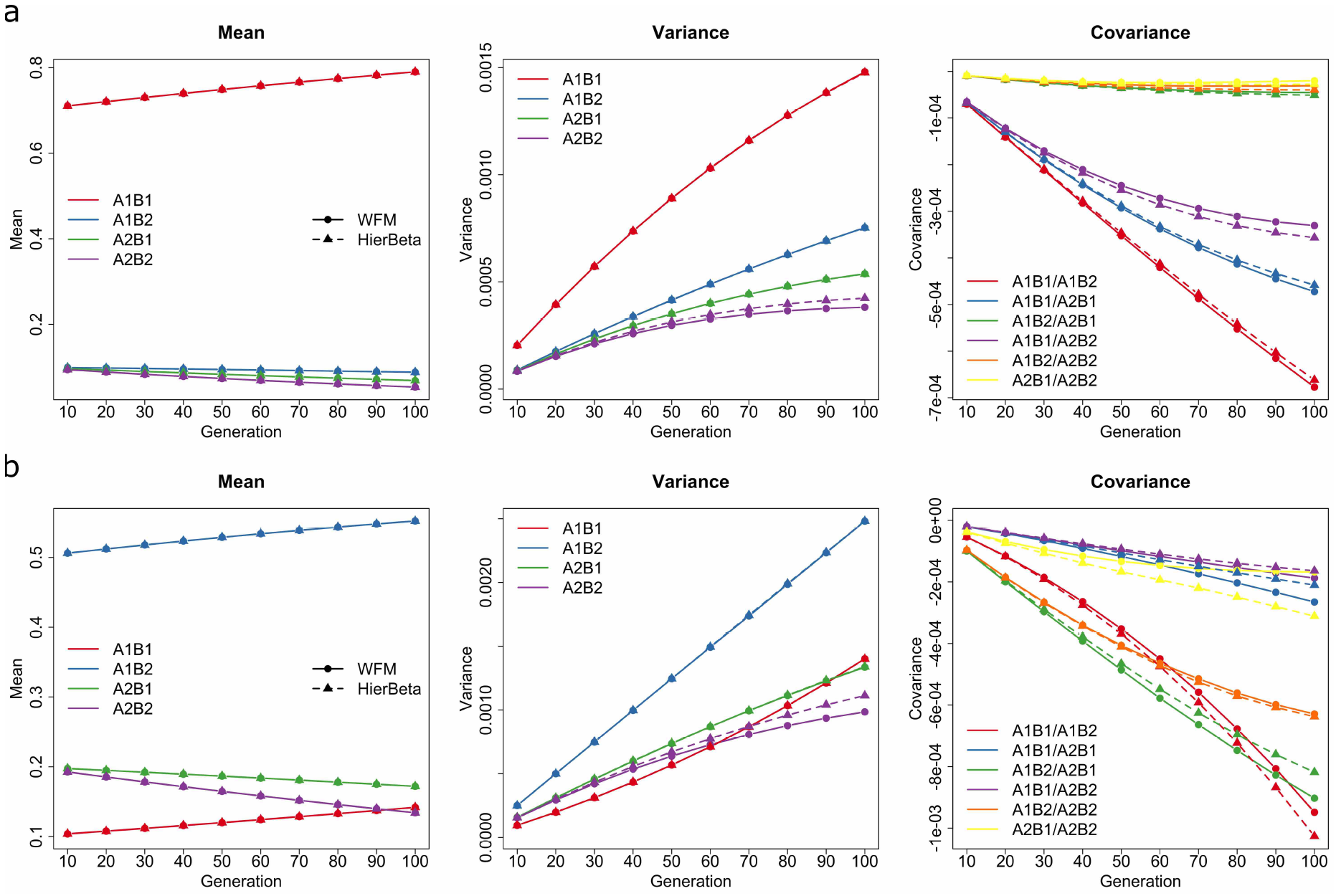
An illustration for the mean, variance and covariance of the Wright-Fisher model and our hierarchical beta approximation, where we set the selection coefficients 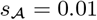 and 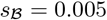, the dominance parameters 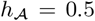 and 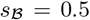, the recombination rate *r* = 0.00001 and the population size *N* = 5000. We pick the initial gamete frequencies (a) ***X***_0_ = (0.7, 0.1, 0.1, 0.1) and (b) ***X***_0_ = (0.1, 0.5, 0.2, 0.2), respectively. The first two moments of the Wright-Fisher model are estimated from 1,000,000 replicated simulations.

Using the law of total covariance, we can write down the covariance between four gamete frequencies under our hierarchical beta approximation in terms of the mean *m*_*i,k*_ and the variance *V*_*i,k*_ of each beta distribution. For example, the covariance between the frequencies of the 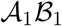 gamete and the 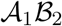 gamete under our hierarchical beta approximation can be represented as

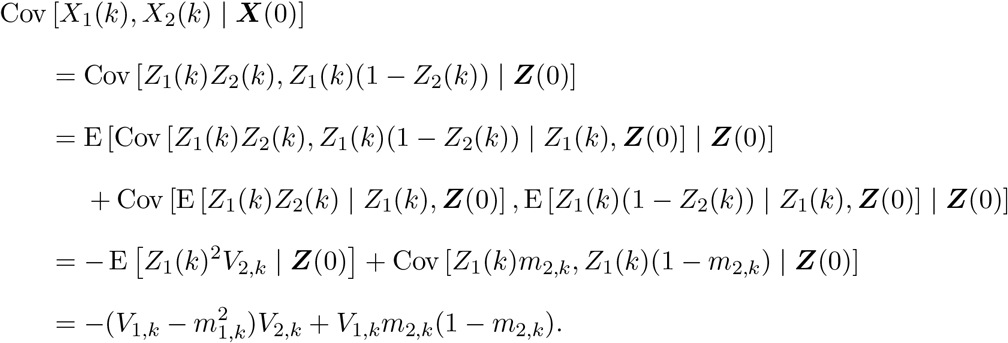

Similarly, we can write down the other covariance between four gamete frequencies under our hierarchical beta approximation as

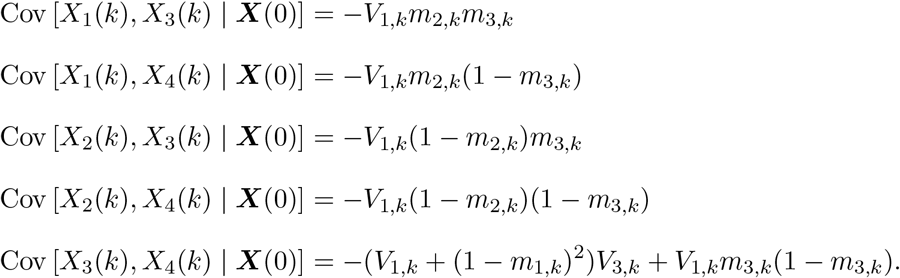

We will discuss how the discrepancy of the (co)variance structure affects the performance of our hierarchical beta approximation through extensive simulations in Section 4.

## 4. Moment-based approximation performance

We evaluate the performance of both our logistic normal and hierarchical beta approximations for the two-locus Wright-Fisher model with selection through extensive simulation studies, comparing them to the diffusion and normal approximations. In what follows, we fix the selection coefficient 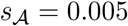 and vary the second selection coefficient 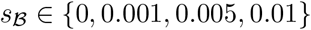, recombination rate *r* ∈ {0.00000001, 0.00001, 0.01} and population size *N* ∈ {500, 5000, 50000}. We take dominance parameters 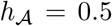 and 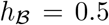 (*i.e.*, the heterozygous fitness is the arithmetic average of the homozygous fitness, called genic selection) unless otherwise noted. In principle, the conclusions hold for any other values of the dominance parameters 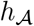 and 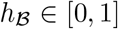. All approximations and simulations described in this work were implemented in R with C++ through the Rcpp and RcppArmadillo packages, and the source code is available at https://github.com/zhangyi-he/WFM-2L-Approx.

### 4.1. Performance of mean and (co)variance approximations

We perform 300 replicated runs for each of the 36 possible combinations of the selection coefficient, recombination rate and population size. For each run, we randomly select the initial gamete frequencies ***X***_0_ from a flat Dirichlet distribution on the state space Ω_***X***_ and simulate 1,000,000 gamete frequency trajectories for 100 generations under the Wright-Fisher model. We compute the empirical mean and (co)variance of the Wright-Fisher model from these 1,000,000 replicates every 10 generations. To find out which method gives the best moment approximation, we approximate the mean and (co)variance every 10 generations using the approach of Terhorst et al. (2015) and our extensions of Lacerda & Seoighe (2014) and Paris et al. (2019). In the sequel, we refer to our three approximation methods as follows:

- Method EL: a scheme using the approximation of mean and (co)variance by our extension of Lacerda & Seoighe (2014).
- Method EP1: a scheme using the approximation of mean and (co)variance by our extension of Paris et al. (2019) up to first order.
- Method EP2: a scheme using the approximation of mean and (co)variance by our extension of Paris et al. (2019) up to second order.

We use the Euclidean vector norm to measure the approximation error of the mean vector, *i.e.*, the difference between the mean under the Wright-Fisher model and their approximations, and the Frobenius matrix norm to measure the approximation error of the (co)variance matrix.

For the case of the population size *N* = 5000, we plot the approximation errors of the mean vector in Figure 3 and the approximation errors of the (co)variance matrix in Figure 4. From Figures 3 and 4, we observe that the error of our approximations increases, *i.e.*, the larger mean error with the larger error bar, with the increase in the selection coefficient and/or recombination rate, and also accumulates with time. A similar conclusion can be obtained for other different population sizes. See Supplemental Material, Figures S1-S4 for the comparison between the mean vector and (co)variance matrix of the Wright-Fisher model and their approximations for the cases of the population size *N* = 500 and *N* = 50000.

**Figure 3:**
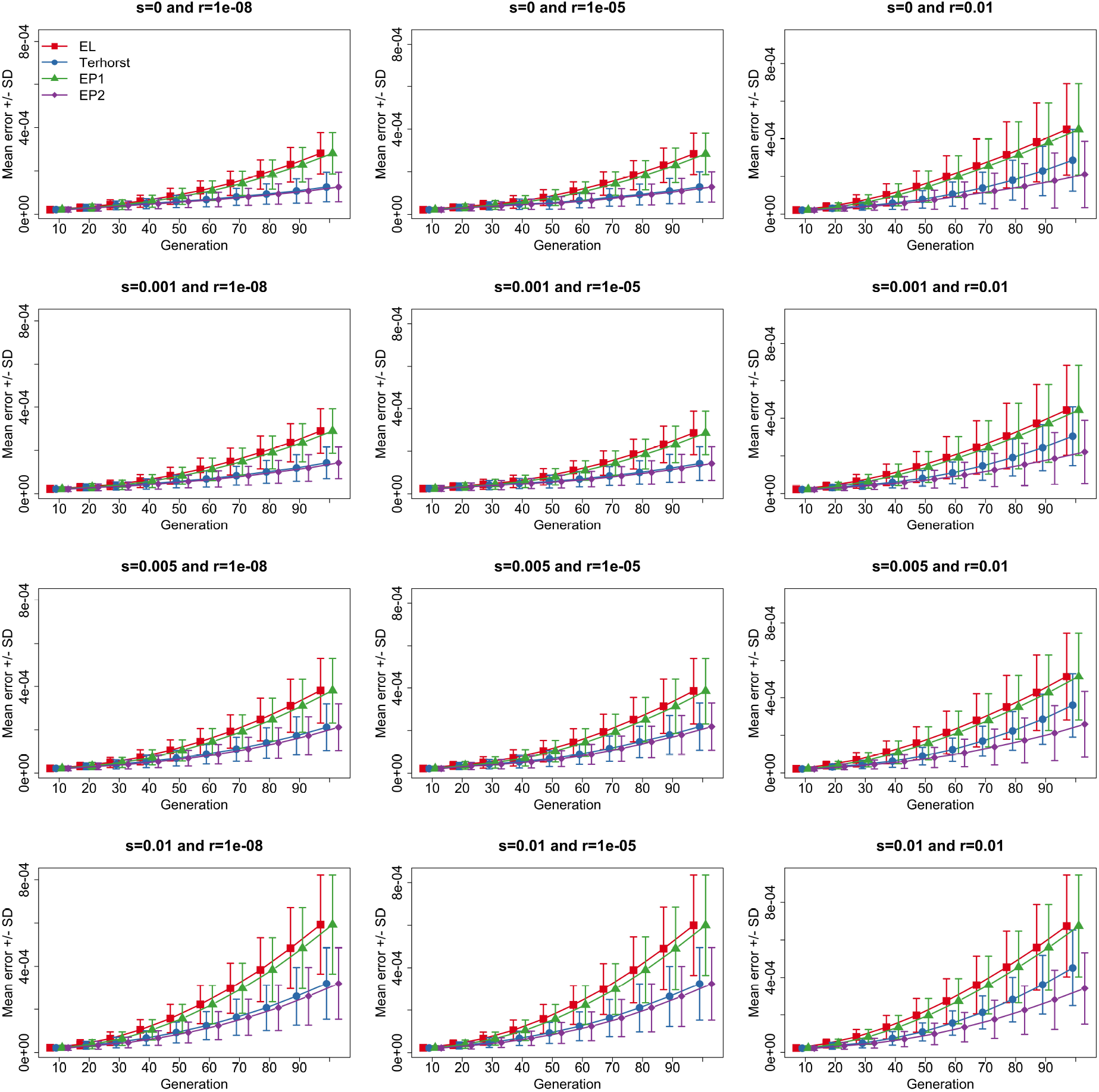
Comparison between the mean vector of the Wright-Fisher model and their approximations for the case of the population size *N* = 5000, where we fix the selection coefficient 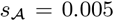 and vary the selection coefficient 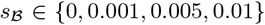 and the recombination rate *r* ∈ {0.00000001, 0.00001, 0.01}. To aid visual comparison of different approximation schemes, we plot the approximation error for each scheme using slightly different *x* coordinate ticks even though the *x* coordinates are exactly the same. We adopt similar practice for all subsequent figures.

**Figure 4:**
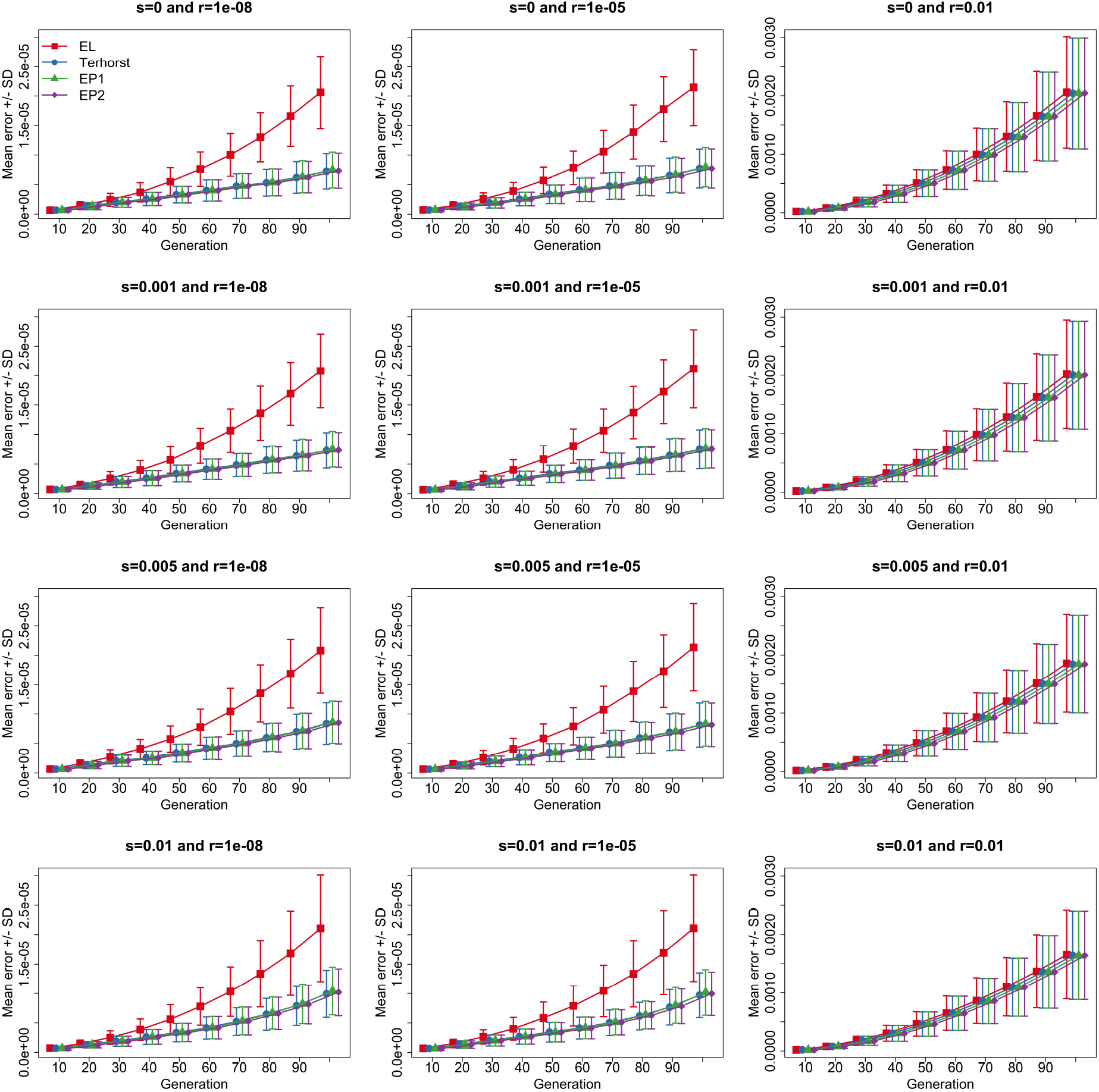
Comparison between the (co)variance matrix of the Wright-Fisher model and their approximations for the case of the population size *N* = 5000, where we fix the selection coefficient 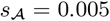 and vary the selection coefficient 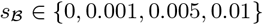 and the recombination rate *r* ∈ {0.00000001, 0.00001, 0.01}.

As illustrated in Figure 3, we get the same performance with methods EL and EP1. Compared to these two methods, the approach of Terhorst et al. (2015) and our method EP2 have better performance across all 12 possible combinations of the selection coefficient and recombination rate. However, the performance advantage of the latter two methods narrows with the increase in the selection coefficient and/or recombination rate. More specifically, our method EP2 matches the performance of the approach of Terhorst et al. (2015) for tightly linked loci (see the left two panels of Figure 3) and becomes better for loosely linked loci (see the right panel of Figure 3). Although the error of all mean approximations increases over time, the approach of Terhorst et al. (2015) sees a much faster increase in the approximation error than the other three methods when the two loci are loosely linked. In Supplemental Material, Figures S1 and S3, we illustrate the results for the cases of the population size *N* = 500 and *N* = 50000. They reveal similar behaviour in approximation errors.

For the approximation error of the (co)variance matrix, we observe from Figure 4 that the approach of Terhorst et al. (2015) and our methods EP1 and EP2 have better performance than our method EL. This advantage is diminished with the decrease in the selection coefficient and/or recombination rate, to nearly non-existent for loosely linked loci (see the right panel of Figure 4). Compared to the approach of Terhorst et al. (2015), our method EP2 matches its performance, but our method EP1 performs slightly worse. Combining with their performance for the cases of population sizes *N* = 500 and *N* = 50000 (see Supplemental Material, Figures S2 and S4), we find that the performance advantage of the approximation of Terhorst et al. (2015) and our methods EP1 and EP2 weakens with the increase in the population size.

In conclusion, moment approximations overall work well. Their performance becomes better with the decrease in the selection coefficient and/or recombination rate, but the approximation error increases with time. Amongst the four moment approximation schemes, our extension of Paris et al. (2019) up to the second order (method EP2) achieves the best performance across all 36 possible combinations of the selection coefficient, recombination rate and population size. More accurate approximations for the mean and (co)variance of the Wright-Fisher model are available if we keep third-or even higher-order terms in the Taylor expansion in our extension of Paris et al. (2019).

### 4.2. Performance of transition probability density approximations

To evaluate the performance of our logistic normal and hierarchical beta approximations for the Wright-Fisher model, we simulate them alongside the diffusion and normal approximations of the Wright-Fisher model, so that we can compare the approximation schemes we introduce with these two widely used methods. More specifically, we perform 300 replicated runs for each of the 36 possible combinations of the selection coefficient, recombination rate and population size. For each run, we randomly draw the starting gamete frequencies ***X***_0_ from a flat Dirichlet distribution over the state space Ω_***X***_ and simulate 1,000,000 gamete frequency trajectories for 100 generations according to the Wright-Fisher model and its approximations. The mean vector and (co)variance matrix of the Wright-Fisher model required in the moment-based approximation are calculated with our extension of Paris et al. (2019) up to the second order, *i.e.*, method EP2 from Section 4.1. The diffusion approximation is simulated with the procedure of He et al. (2020a). Evaluation of the quality of the approximations is carried out by comparing the empirical cumulative distribution functions (CDFs) computed from these simulated gamete frequency trajectories. We randomly choose 5000 vectors of gamete frequencies from a flat Dirichlet distribution on the state space Ω_***X***_, and the empirical CDF is evaluated at these sampled gamete frequencies at every 10 generations from the 1,000,000 simulated gamete frequency trajectories. We measure the difference between the Wright-Fisher model and its approximation using the root mean squared difference (RMSD) between the two empirical CDFs.

We summarise the RMSD between the Wright-Fisher model and its approximations in Figures 5–7 for all 36 possible combinations of the selection coefficient, recombination rate and population size. Compared to the three moment-based approximations, the diffusion approximation shows better overall performance across all 36 possible combinations in terms of computational accuracy, *i.e.*, the lowest mean error with the smallest error bar, even though some combinations of the selection coefficient, recombination rate and population size do not completely satisfy the assumption required in the diffusion approximation (see, *e.g.*, He et al., 2020a). However, in terms of computational efficiency, the diffusion approximation performs significantly worse than the moment-based approximations. For example, to construct the empirical CDFs at every 10 generations in the top left plot of Figures 5, we generate 1,000,000 realisations. This takes about 181.358 seconds on a single core of an Intel Core i7 processor at 4.2 GHz with the diffusion approximation. In contrast, this only takes approximately 0.001 seconds with the moment-based approximations. Additionally, using diffusion approximation, at every 10 generations 1,000,000 realisations are required to be stored for the computation of the empirical CDFs, 341.386 MB in total, whereas in moment-based approximations we only need to store the relevant parameters, *e.g.*, the shape parameters for the hierarchical beta distribution at every 10 generations, 523.2 bytes in total.

**Figure 5:**
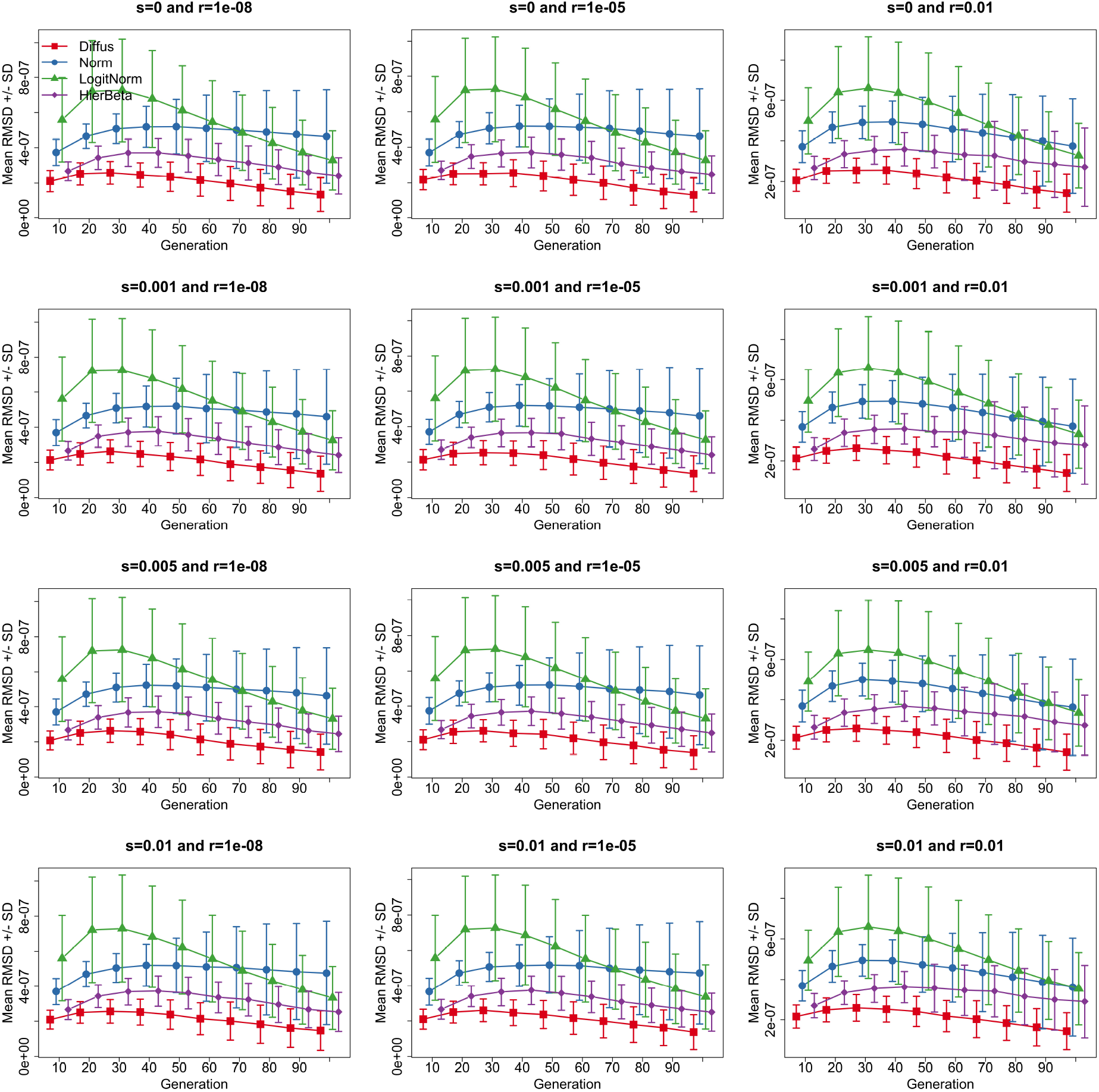
Comparison between the empirical CDFs of the Wright-Fisher model and their approximations for the case of the population size *N* = 500, where we fix the selection coefficient 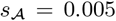 and vary the selection coefficient 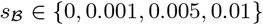 and the recombination rate *r* ∈ {0.00000001, 0.00001, 0.01}.

**Figure 6:**
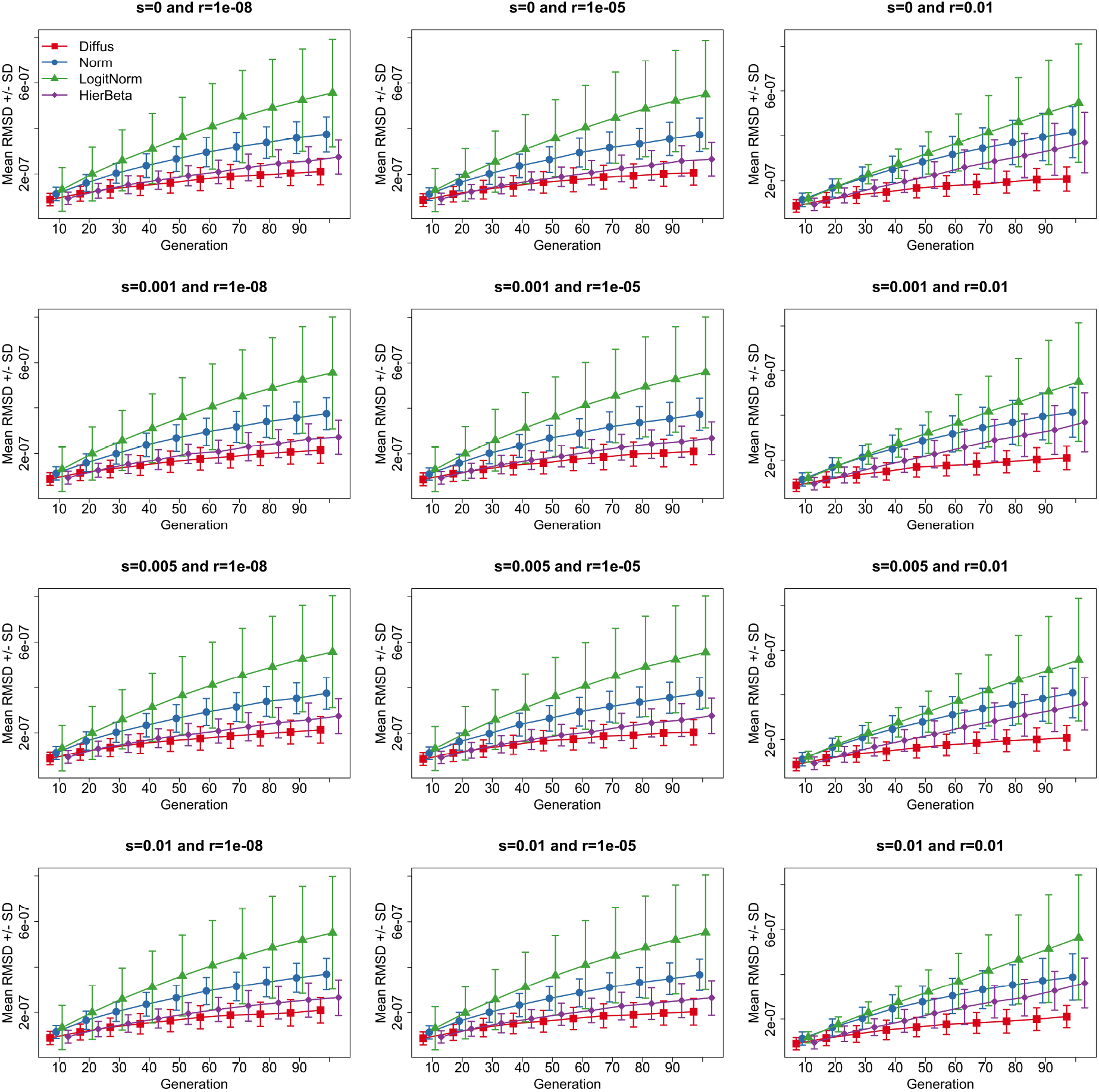
Comparison between the empirical CDFs of the Wright-Fisher model and their approximations for the case of the population size *N* = 5000, where we fix the selection coefficient 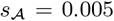 and vary the selection coefficient 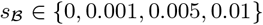 and the recombination rate *r* ∈ {0.00000001, 0.00001, 0.01}.

**Figure 7:**
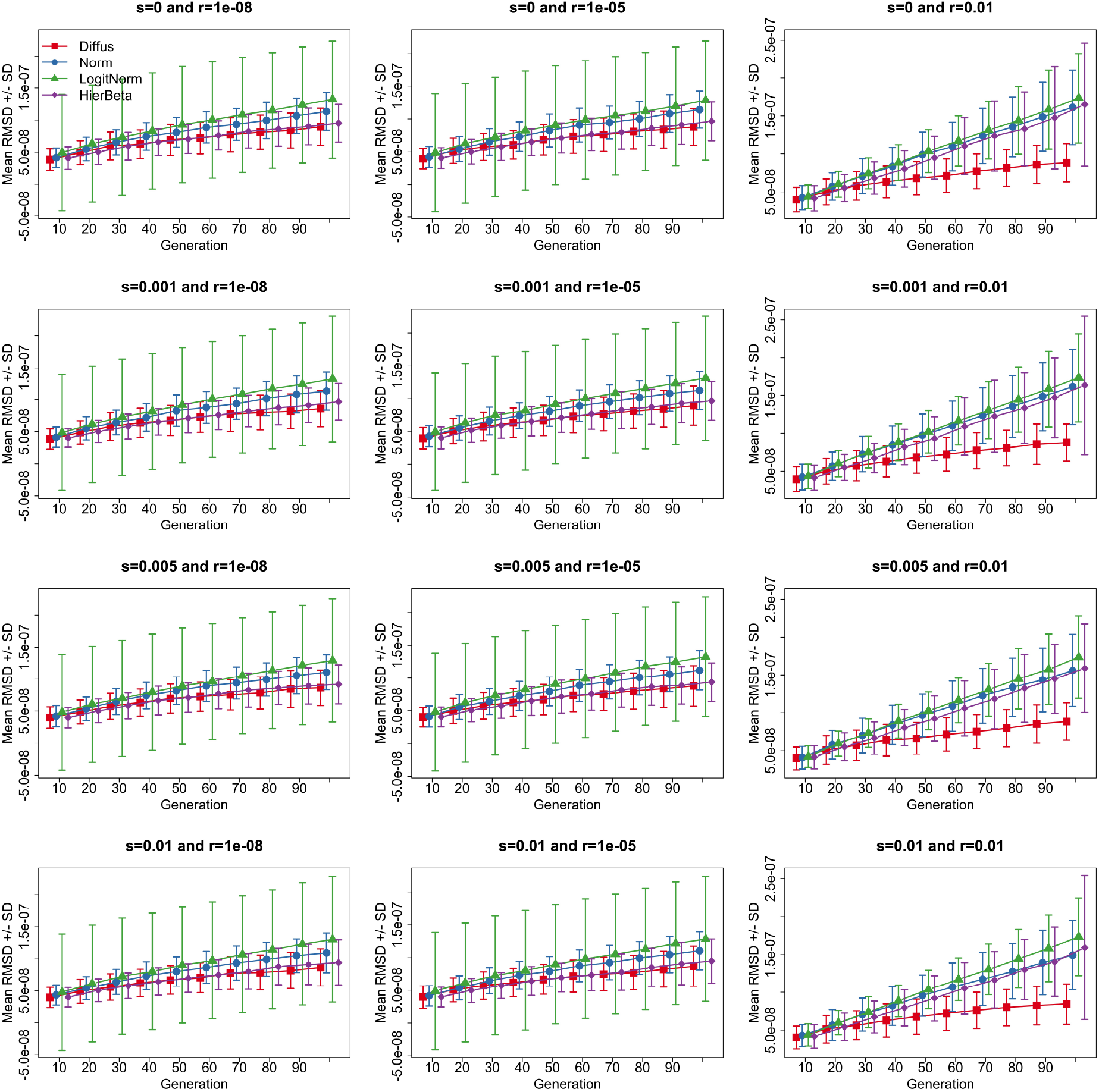
Comparison between the empirical CDFs of the Wright-Fisher model and their approximations for the case of the population size *N* = 50000, where we fix the selection coefficient 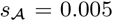 and vary the selection coefficient 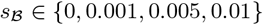 and the recombination rate *r* ∈ {0.00000001, 0.00001, 0.01}.

As illustrated in Figures 5–7, moment-based approximations have better overall performance as the population size increases and become worse with longer times. This is consistent with the conclusion for the single-locus case presented in Lacerda & Seoighe (2014). In the three moment-based approximations, our logistic normal approximation achieves the worst overall performance except for small population sizes. We find from Figure 5 that our logistic normal approximation achieves the worst performance during the initial tens of generations and then achieves better performance than the normal approximation with longer times for small population sizes. In general, alleles drift toward loss or fixation much faster in smaller populations, which will result in a leak of probability mass into the values of the gamete frequencies that are out of the state space Ω_***X***_ owing to the infinite support of the normal distribution. Therefore, our logistic normal approximation works better than the normal approximation only when the allele frequencies approach the boundaries.

Figures 5–7 also show that our hierarchical beta approximation yields the best overall performance amongst the three moment-based approximations across all 36 possible combinations of the selection coefficient, recombination rate and population size and even matches the performance of the diffusion approximation for small recombination rates and large population sizes (see the left two panels of Figure 7). However, it should be noted that in practice non-positive shape parameters will be produced in our hierarchical beta approximation from some mean and (co)variance of the Wright-Fisher model, which is not mathematically possible. We summarise the proportion of the cases that the Wright-Fisher model cannot be matched by our hierarchical beta approximation at every 10 generations for each possible combination in Figure 8. We observe that the failure proportion has a trend of slow increase over time for small recombination rates but shows a sharp increase for large recombination rates. We also see from Figure 8 that the failure proportion slightly increases with the decrease in the population size, which becomes significant for large recombination rates. Such a computational instability issue could be caused by that our moment approximation gets less accurate with time and our hierarchical beta approximation fails to model fixation or loss of alleles.

**Figure 8:**
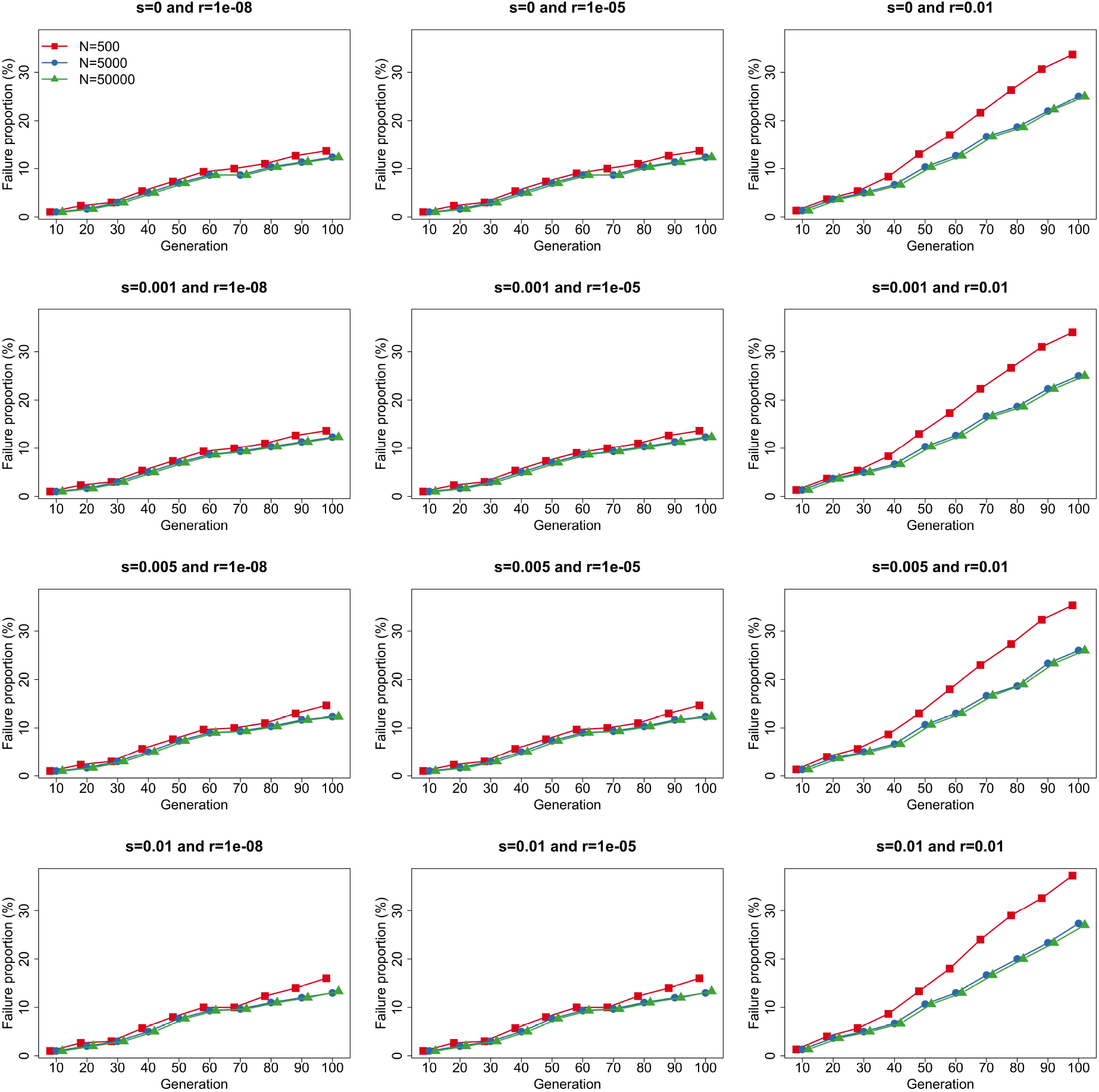
Proportions of the cases that the Wright-Fisher model cannot be fitted through our hierarchical beta approximation across all 36 possible combinations of the selection coefficient, recombination rate and population size.

To resolve the computational instability issue of our hierarchical beta approximation, if the three selected gamete frequencies, *e.g.*, *X*_1_(*k*), *X*_2_(*k*) and *X*_3_(*k*), fail to produce positive shape parameters for each beta distribution, we choose other three gamete frequencies to construct our hierarchical beta approximation instead. Permuting the indices of our hierarchical beta approximation can completely resolve the failure cases for small recombination rates (see the left two panels of Figure 9) and significantly reduce the failure proportions for large recombination rates (see the right panel of Figure 9). For the remaining cases of failure, *i.e.*, those which permutation of the indices of the hierarchical beta approximation cannot resolve, we search for nearby mean and variance values that do yield hierarchical beta approximation with positive shape parameters. Such a procedure completely eliminates the computational instability issue but would result in additional errors. In the simulation studies we have performed, we have used the procedure outlined in this paragraph. As shown in Figures 5–7, our hierarchical beta approximation achieves the best overall performance among the three moment-based approximations even though it needs these failure prevention steps.

**Figure 9:**
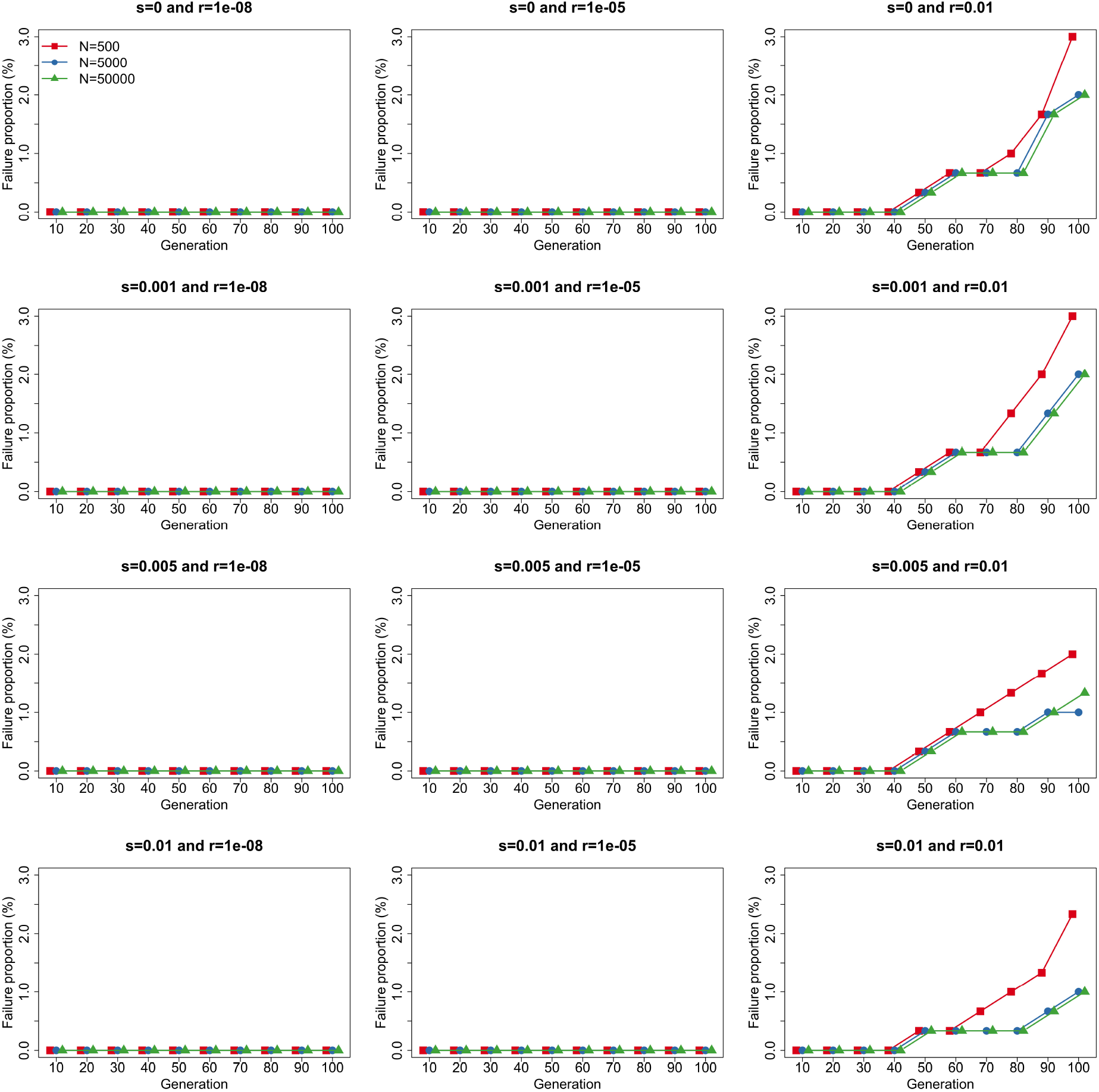
Proportions of the cases that the Wright-Fisher model cannot be fitted through our hierarchical beta approximation (with permutation) across all 36 possible combinations of the selection coefficient, recombination rate and population size. The remaining failure cases are resolved by the failure prevention steps as explained above.

In summary, the moment-based approximations enable us to compute the approximate transition probabilities of the Wright-Fisher model at a much smaller computational cost than the diffusion approximation, even though the diffusion approximation achieves higher computational accuracy. Compared to the normal approximation of Terhorst et al. (2015), our logistic normal approximation achieves higher computational accuracy only when the Wright-Fisher model gets close to the boundaries. Our hierarchical beta approximation achieves the highest computational accuracy among the three moment-based approximations and even matches the performance of the diffusion approximation for small recombination rates and large population sizes. For large recombination rates, the performance advantage of our hierarchical beta approximation is diminished. However, the two loosely linked loci are approximately independent with each other due to the large number of recombinations, therefore can typically be characterised through the single-locus Wright-Fisher model in practice.

## 5. Discussion

In this work, we proposed two moment-based approximations for the two-locus Wright-Fisher model with selection, the logistic normal approximation and the hierarchical beta approximation. The main idea is to match the true moments of the Wright-Fisher model with the fitted moments of a parametric family of probability distributions, dating back to Cavalli-Sforza & Edwards (1967). A prerequisite for the moment-based approximation is that we can calculate the mean and (co)variance of the two-locus Wright-Fisher model with selection at any given time point, so we extended the approaches of Lacerda & Seoighe (2014) and Paris et al. (2019), which in their original form were limited to the Wright-Fisher model of population dynamics under natural selection at a single locus. A major advantage of our logistic normal and hierarchical beta approximations, compared to *e.g.*, the Gaussian approximation of Terhorst et al. (2015), is that we avoid the distribution support issue.

Through extensive simulations studies, we illustrated that our approximations for the mean and (co)variance achieved better performance with smaller selection coefficients and/or recombination rates, but has deteriorating performance with the increase of time horizon, owing to the accumulation of approximation errors. Our extension of Paris et al. (2019) up to the second order achieved the best performance in the four moment approximations we compared. In our simulation studies, our moment-based approximations could not yield the same computational accuracy as the diffusion approximation but enabled the computation of the transition probabilities of the Wright-Fisher model at a fraction of the computational cost. The performance of the moment-based approximations was better for larger population sizes but deteriorated with the increase of time horizon. This is consistent with the conclusion stated in Lacerda & Seoighe (2014) for the single-locus case. The performance of our logistic normal approximation was not as good as the Gaussian approximation of Terhorst et al. (2015) except when the Wright-Fisher model approached the boundaries and began to be absorbed. In contrast, our hierarchical beta approximation outperformed the Gaussian approximation and even matched the performance of the diffusion approximation for small recombination rates and large population sizes.

In recent years, the Wright-Fisher model has been successfully applied to the analysis of time series allele frequency data. However, most of these methods are limited to either a single locus or multiple independent loci, *i.e.*, genetic recombination effect and local linkage information are ignored in these methods. This may lead to a poor inference of natural selection, especially for the case of tightly linked loci (He et al., 2020b). Existing approaches that allow us to account for genetic recombination and local linkage suffer from either distribution support issues in Terhorst et al. (2015) or prohibitive computational costs in He et al. (2020b). Our moment-based approximations provide an alternative way to carry out the likelihood computation. For example, our hierarchical beta approximation has been shown to balance computational accuracy and efficiency in our simulation studies. It can therefore be expected to improve the performance of existing statistical frameworks for the inference of natural selection from time series genetic data. Moreover, our hierarchical beta approximation makes it possible to conduct genome-wide inference of natural selection from time series genomic data, modelling genetic recombination and local linkage in a maximum composite likelihood framework. Since the computational accuracy of our hierarchical beta approximation becomes worse with the increase of time horizon, it is better suited to the analysis of the genetic data of short evolutionary time scales such as experimental evolution studies (*e.g.*, Lang et al., 2013; Wiser et al., 2013; Burke et al., 2014; Le Bihan-Duval et al., 2018; Papkou et al., 2019; Barghi et al., 2019).

A number of improvements to our moment-based approximations may be possible. First of all, our moment-based approximations cannot completely capture the (co)variance structure of the two-locus Wright-Fisher model with selection although their performance has been shown to be acceptable. Recently, Speed et al. (2019) developed the pyramid hierarchical beta approximation of the Wright-Fisher model for the evolution of relative frequencies of four alleles at a single locus, which has enough free parameters to characterise the entire (co)variance structure. However, it cannot be extended to the two-locus Wright-Fisher model with selection since only a small subset of the state space can guarantee the existence of positive shape parameters for each beta distribution in their pyramid hierarchical beta approximation. It would be fruitful to find other parametric families of probability distributions that enable the complete capture of the (co)variance structure of the two-locus Wright-Fisher model with selection. Second, our moment-based approximations avoid the distribution support issue found in Terhorst et al. (2015) but fail to model fixation or loss of alleles. Tataru et al. (2015) addressed this issue and introduced the beta-with-spikes approximation for the Wright-Fisher model for the evolution of relative frequencies of two alleles at a single locus. Paris et al. (2019) extended it to incorporate natural selection and applied it to the inference of natural selection from time series data of allele frequencies. In their work, modelling fixation or loss of alleles has been shown to improve the performance of the beta approximation. It would be interesting to find ways to capture the behaviour of the two-locus Wright-Fisher model with selection at the boundaries.

## Acknowledgements

This work was funded in part by the Engineering and Physical Sciences Research Council (EPSRC) Grant EP/I028498/1 to F.Y.

## Appendix A. Derivation of the moment approximations

We demonstrate how to extend the moment approximations developed in Lacerda & Seoighe (2014) and Paris et al. (2019) to the two-locus Wright-Fisher model with selection in this section, including the first- and second-order approximations proposed by Paris et al. (2019). We also introduce the moment approximations developed in Terhorst et al. (2015).

## Appendix A.1. Extension of Lacerda & Seoighe (2014)

In this section, we provide the detailed derivation of the extension of the moment approximation proposed in Lacerda & Seoighe (2014) to the two-locus Wright-Fisher model with selection. Using the law of total mean and (co)variance, we can formulate the mean and (co)variance of the gamete frequencies ***X***(*k*) given the gamete frequencies ***X***(0) as

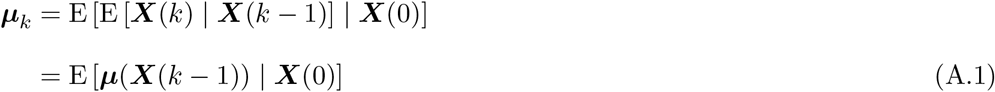

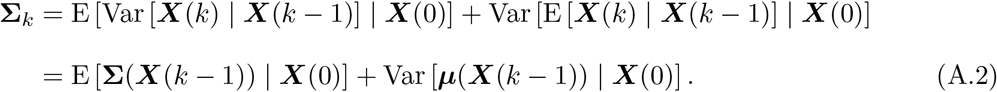

Following Lacerda & Seoighe (2014), we approximate the three terms on the right-hand side of Eqs. (A.1) and (A.2) with the multivariate delta method (see Oehlert, 1992, for more details), thereby obtaining

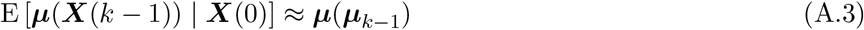

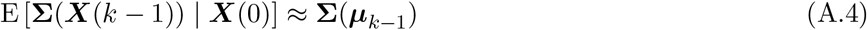

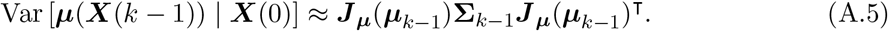

Substituting Eqs. (A.3)–(A.5) into Eqs. (A.1) and (A.2), we have

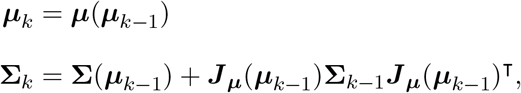

which is the recursion formula for the approximation of the mean and (co)variance of the two-locus Wright-Fisher model with selection extended from Lacerda & Seoighe (2014). The initial mean and (co)variance are ***μ***_0_ = E[***X***(0)] and **Σ**_0_ = Var[***X***(0)], respectively.

## Appendix A.2. Extension of Paris et al. (2019)

In this section, we show how to extend the moment approximation of Paris et al. (2019) to the two-locus Wright-Fisher model with selection. Following Paris et al. (2019), we substitute Eqs. (4) and (5) into Eq. (7), and then the (co)variance of the gamete frequencies ***X***(*k*) given the gamete frequencies ***X***(0) can be formulated as

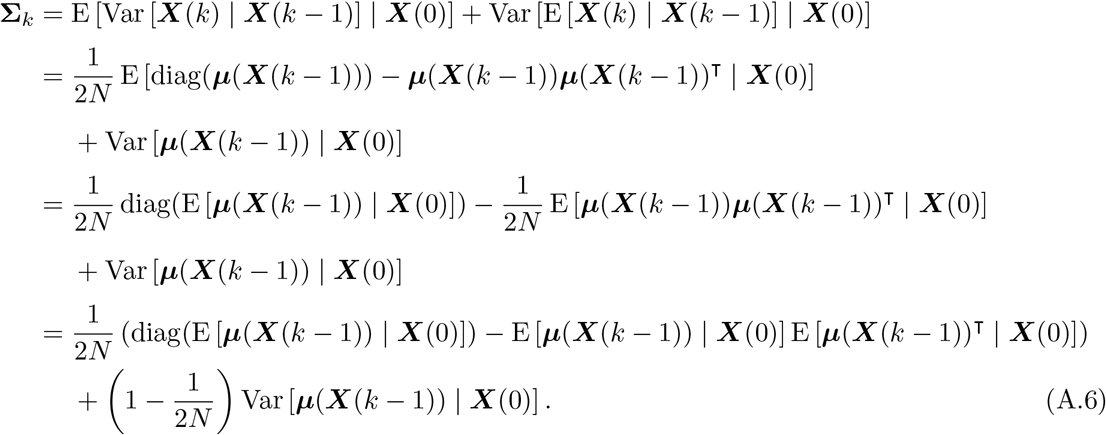

Using the Taylor expansion of ***μ***(***X***(*k* − 1)) about ***μ***_*k*−1_ up to the first order, we have

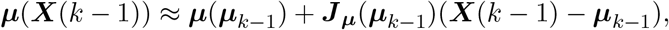

thereby obtaining

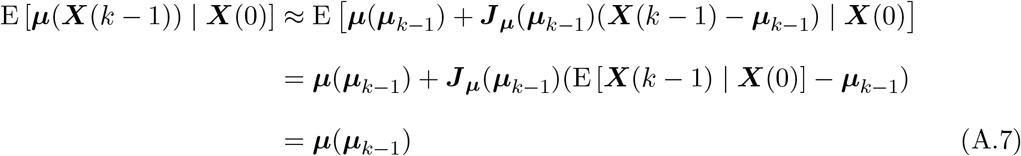

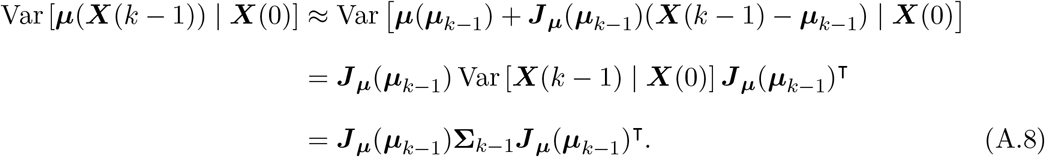

Substituting Eqs. (A.7) and (A.8) into Eqs. (A.1) and (A.6), we have

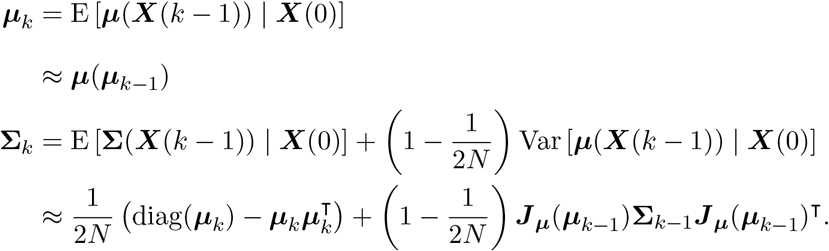

Using Eqs. (4) and (5), we can formulate the recursion formula for the first-order approximation of the mean and (co)variance of the two-locus Wright-Fisher model with selection extended from Paris et al. (2019) as

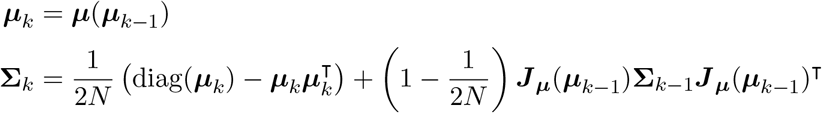

with the initial values ***μ***_0_ = E[***X***(0)] and **Σ**_0_ = Var[***X***(0)].

Similarly, with the Taylor expansion of ***μ***(***X***(*k* − 1)) about ***μ***_*k−*1_ up to the second order

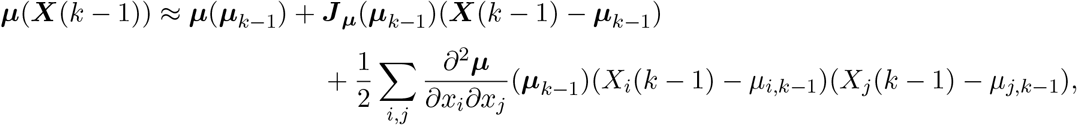

the recursion formula for the second-order approximation of the mean and (co)variance of the two-locus Wright-Fisher model with selection can be written down as

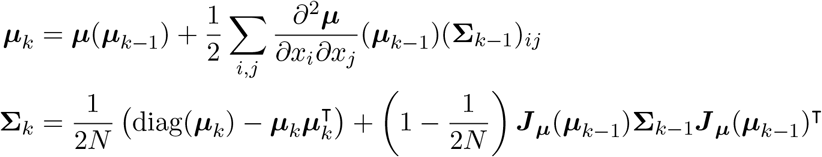

with the initial values ***μ***_0_ = E[***X***(0)] and **Σ**_0_ = Var[***X***(0)].

## Appendix A.3. Introduction of Terhorst et al. (2015)

In this section, we show how to apply the moment approximation of Terhorst et al. (2015) to the two-locus Wright-Fisher model with selection. Following Terhorst et al. (2015), we write the gamete frequencies ***X***(*k*) = ***x**_k_* +*δ**X***(*k*), where ***x***_0_ = E[***X***(0)] and ***x**_k_* = ***μ***(***x***_*k−*1_) for *k* ∈ ℕ^+^, and introduce two correction terms ***ϵ***_*k*_ = E [*δ**X***(*k*) | ***X***(0)] and ***ξ***_*k*_ = E [*δ**X***(*k*)*δ**X***(*k*)^┬^ | ***X***(0)].

Unlike Paris et al. (2019), we use the Taylor expansion of ***μ***(***X***(*k* − 1)) about ***x***_*k−*1_ up to the second order rather than ***μ***_*k−*1_, and then we have

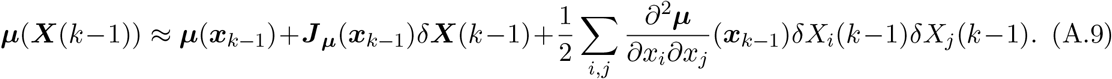

With Eq. (A.1), the mean of the gamete frequencies ***X***(*k*) given the gamete frequencies ***X***(0) can be represented as

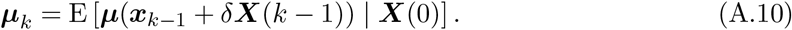

Substituting Eq. (A.9) into (A.10), we have

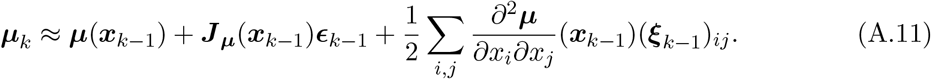

Similarly, with Eq. (A.2), the (co)variance of the gamete frequencies ***X***(*k*) given the gamete frequencies ***X***(0) can be expressed as

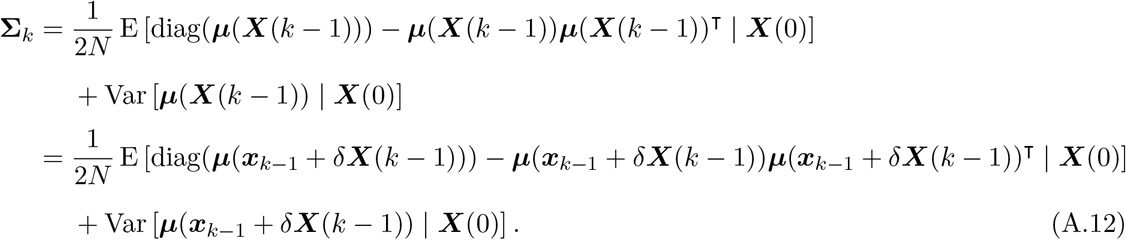

Substituting Eq. (A.9) into (A.12), we have

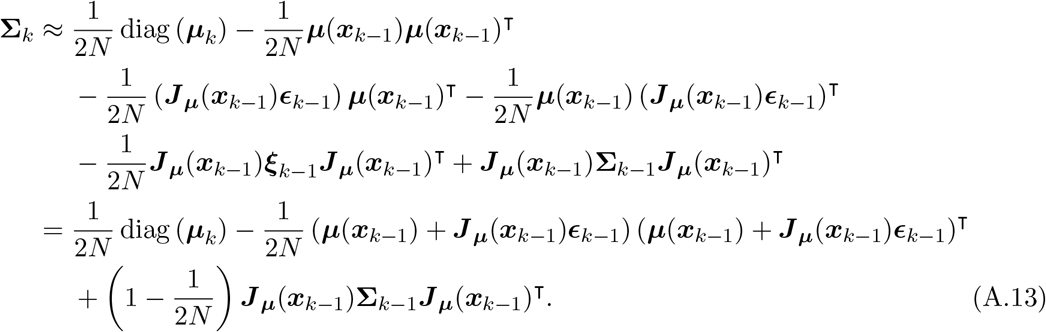

By definition, we can formulate the mean and (co)variance of the gamete frequencies ***X***(*k*) given the gamete frequencies ***X***(0) as

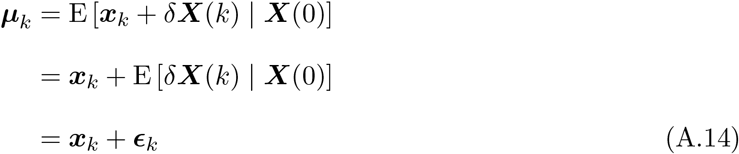

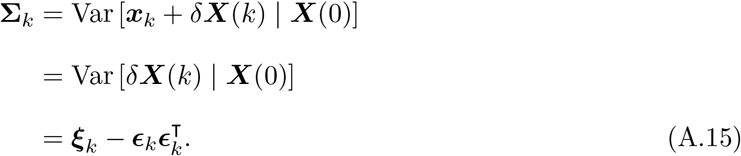

Combining Eqs. (A.11), (A.13), (A.14) and (A.15), we can write down the recursion formula for the approximation of the mean and (co)variance of the two-locus Wright-Fisher model with selection introduced from Terhorst et al. (2015) as

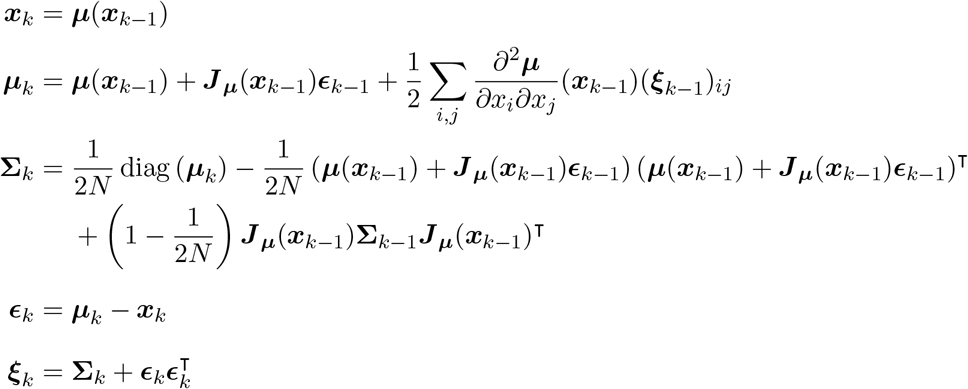

with the initial values *x*_0_ = *μ*_0_ = E[*X*(0)], **Σ**_0_ = Var[*X*(0)], *ϵ*_0_ = **0** and *ξ*_0_ = **0**. Note that **0** denotes the four-dimensional zero vector or the four by four zero matrix as appropriate.

