## Supplemental Material for "Moment-based approximations for the Wright-Fisher model of population dynamics under natural selection at two linked loci"

### File S1. Performance evaluation of moment approximations

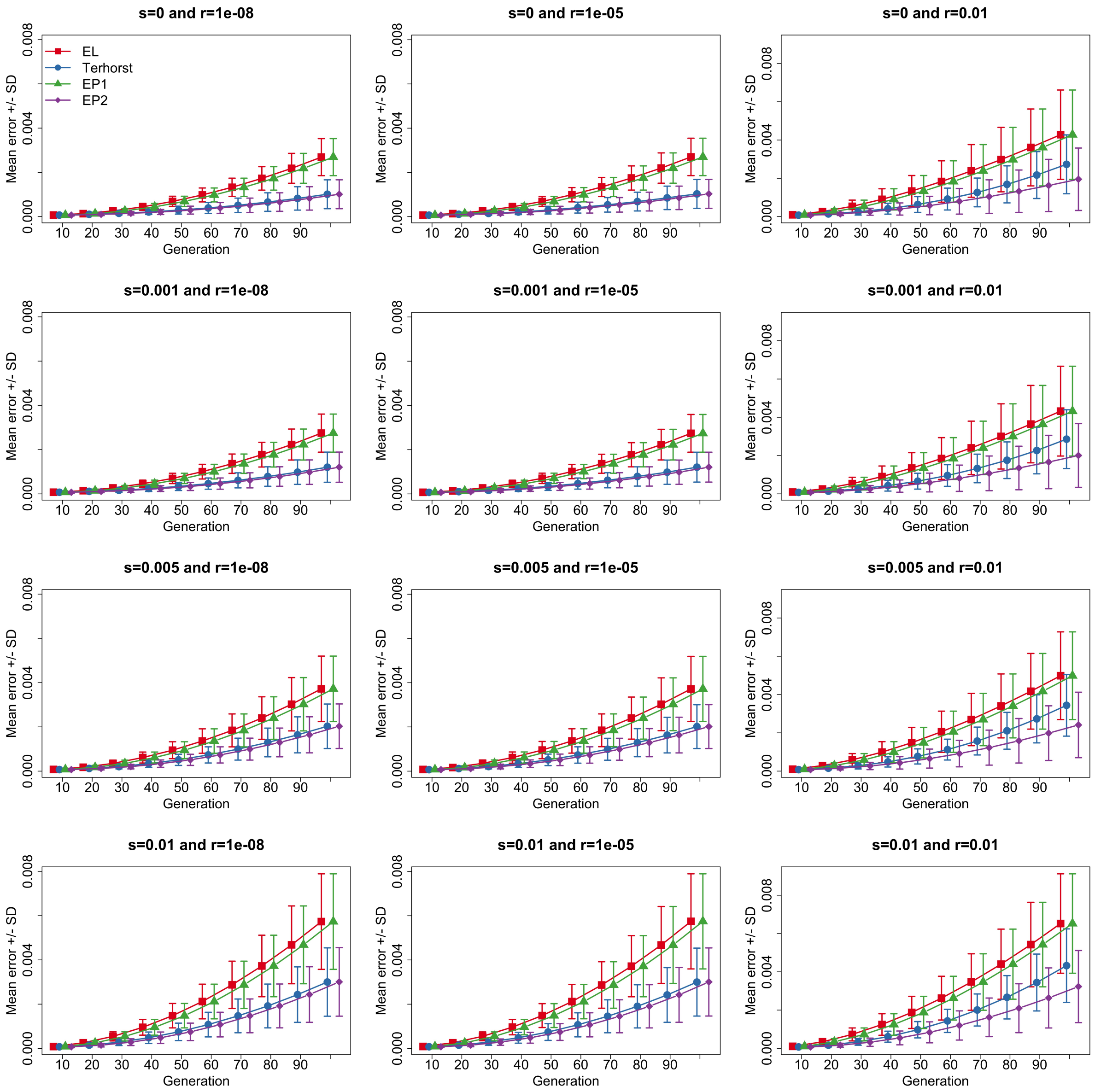


Figure S1: Comparison between the mean vector of the Wright-Fisher model and their approximations for the case of the population size $N = 500$, where we fix the selection coefficient $s_{\mathcal{A}} = 0.005$ and vary the selection coefficient $s_{\mathcal{B}} \in \{0, 0.001, 0.005, 0.01\}$ and the recombination rate $r \in \{0.00000001, 0.00001, 0.01\}$. To aid visual comparison of different approximation schemes, we plot the approximation error for each scheme using slightly different $x$ coordinate ticks even though the $x$ coordinates are exactly the same. We adopt similar practice for all subsequent figures.

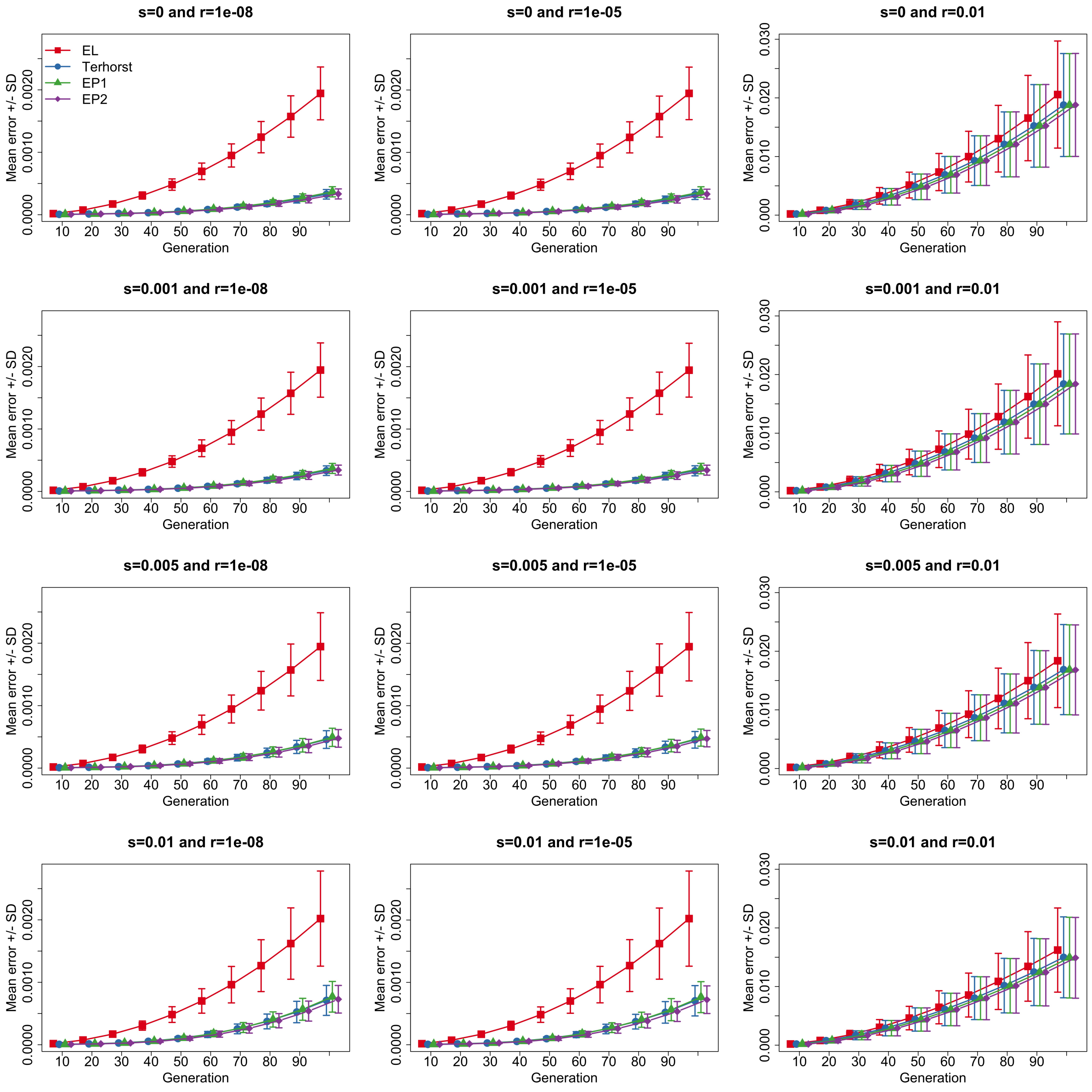


Figure S2: Comparison between the (co)variance matrix of the Wright-Fisher model and their approximations for the case of the population size $N = 500$, where we fix the selection coefficient $s_{\mathcal{A}} = 0.005$ and vary the selection coefficient $s_{\mathcal{B}} \in \{0, 0.001, 0.005, 0.01\}$ and the recombination rate $r \in \{0.00000001, 0.00001, 0.01\}$.

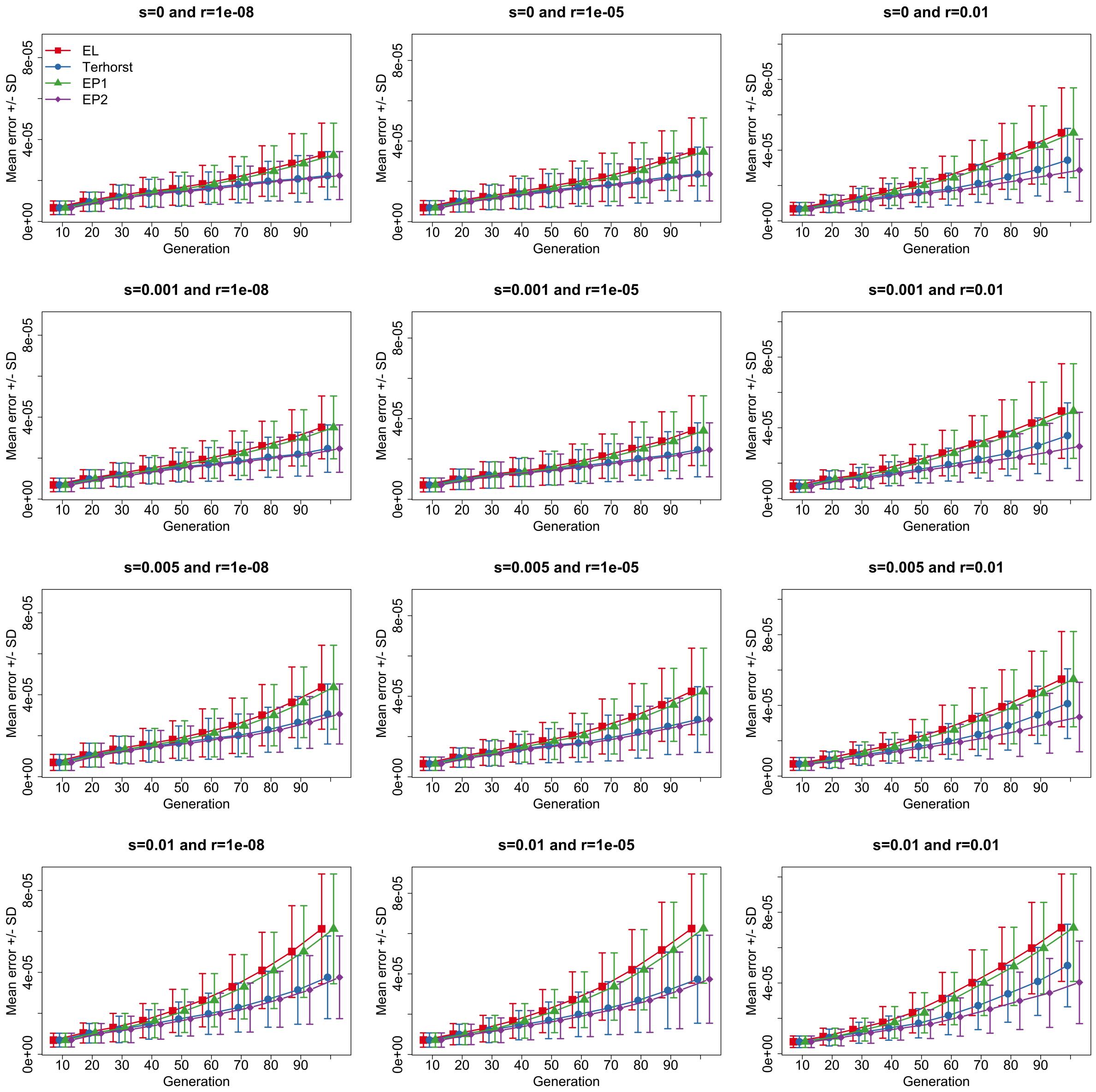


Figure S3: Comparison between the mean vector of the Wright-Fisher model and their approximations for the case of the population size $N = 50000$, where we fix the selection coefficient $s_{\mathcal{A}} = 0.005$ and vary the selection coefficient $s_{\mathcal{B}} \in \{0, 0.001, 0.005, 0.01\}$ and the recombination rate $r \in \{0.00000001, 0.00001, 0.01\}$.

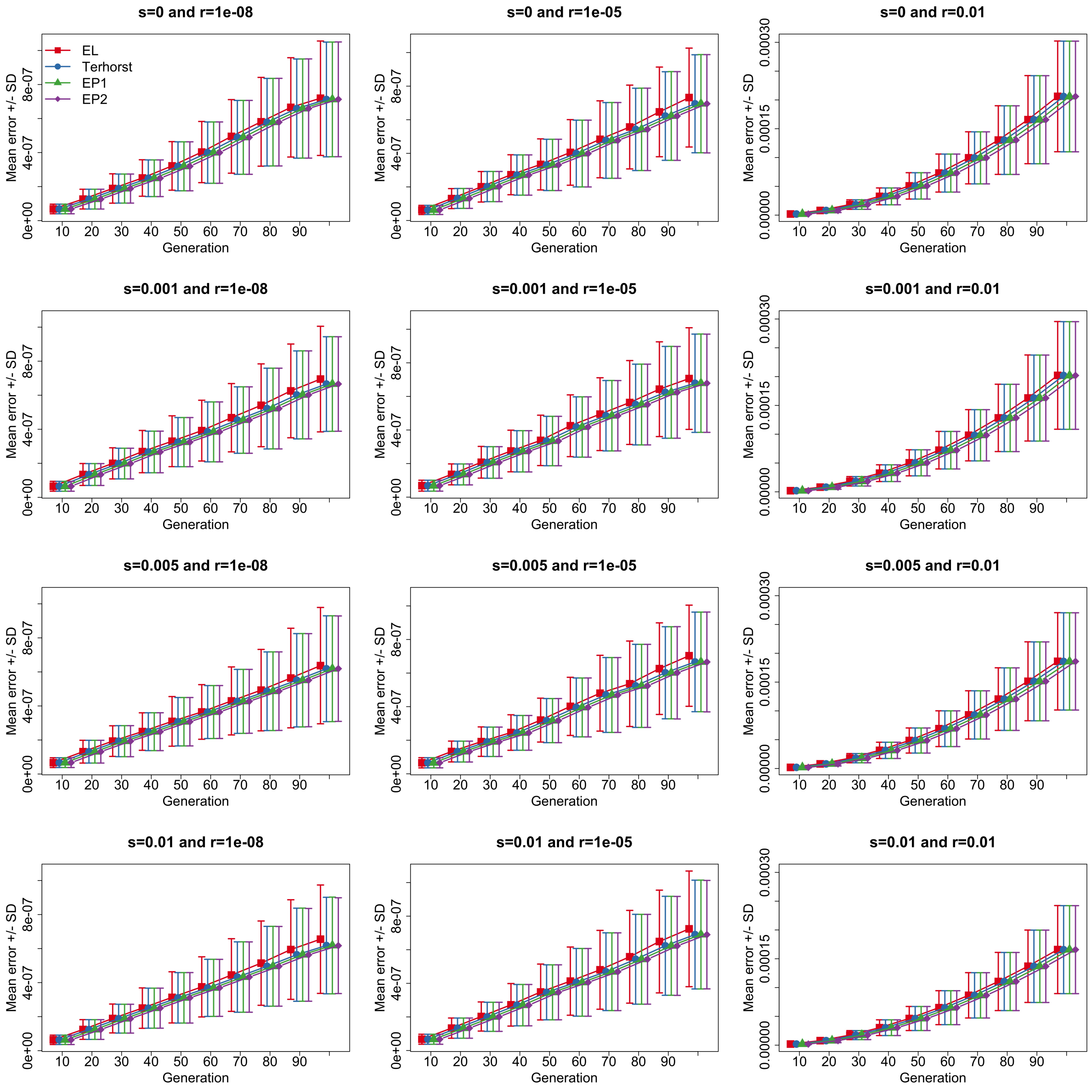


Figure S4: Comparison between the (co)variance matrix of the Wright-Fisher model and their approximations for the case of the population size $N = 50000$, where we fix the selection coefficient $s_{\mathcal{A}} = 0.005$ and vary the selection coefficient $s_{\mathcal{B}} \in \{0, 0.001, 0.005, 0.01\}$ and the recombination rate $r \in \{0.00000001, 0.00001, 0.01\}$.
